# Engineering oncogenic hotspot mutations on *SF3B1* via CRISPR-directed PRECIS mutagenesis

**DOI:** 10.1101/2024.02.23.581842

**Authors:** Mike Fernandez, Qiong Jia, Lei Yu, Xuesong Wang, Kevyn Hart, Zhenyu Jia, Ren-Jang Lin, Lili Wang

## Abstract

*SF3B1* is the most recurrently mutated RNA splicing factor in cancer; However, its study has been hindered by a lack of disease-relevant cell line models. Here, we compared four genome engineering platforms to establish *SF3B1*mutant cell lines: CRISPR-Cas9 editing, AAV HDR editing, base editing (ABEmax, ABE8e), and prime editing (PE2, PE3, PE5Max). We showed that prime editing via PE5max achieved the most efficient *SF3B1* K700E editing across a wide range of cell lines. We further refined our approach by coupling prime editing with a with a fluorescent reporter that leverages a *SF3B1* mutation-responsive synthetic intron to mark prime edited cells. Calling this approach prime editing coupled intron-assisted selection (PRECIS), we then introduced the K700E hotspot mutation into two chronic lymphocytic leukemia (CLL) cell lines, HG-3 and MEC-1, and demonstrated that our PRECIS-engineered cells faithfully recapitulate the altered splicing and copy number variation (CNV) events frequently found in CLL patients with *SF3B1* mutation. Our results showcase PRECIS as an efficient and generalizable method for engineering genetically faithful *SF3B1* mutant models, shed new light on the role of *SF3B1* mutation in cancer biology, and enables generation of novel *SF3B1* mutant cell lines in any cellular context.

## Introduction

Advances in gene editing technologies promise to revolutionize cancer research via precise and accurate modeling of specific point mutations^1^. This is particularly relevant for the study of *SF3B1*, an RNA splicing factors involved in the U2 snRNP recognition of the 3’ splice site (SS) during pre-mRNA splicing, that is recurrently mutated in multiple different leukemias and solid tumors^2^. Hotspot mutations on *SF3B1* drive cryptic 3’ splice site selection, rewiring cellular splicing circuitries and promoting oncogenesis in a variety of cancers^3–7^. Murine models with lineage-restricted expression of *Sf3b1* mutation have enabled mechanistic and functional dissection of the role of *SF3B1 in vivo*^8,9^; However, these models are restricted by high operating cost limiting the availability of different cancer models, human-to-mouse disease heterogeneity, and difficulty in further genetic manipulation. *In vitro* cell line systems are therefore an attractive orthogonal model for the functional and mechanistic dissection of *SF3B1* mutation owing to the ease of engineering and broad representation from across the cancer spectrum.

Based on COSMIC v97 database, *SF3B1* is recurrently mutated in many different types of cancers with a particularly high frequency for hematological malignancies at the K700E position on exon 15^10,11^ (**Fig. 1a-c**). While several isogenic *SF3B1* mutant cell lines have been generated (supplementary table 1)^12–15^, the majority of these do not model *SF3B1* mutation in the proper cancer contexts. For example, while *SF3B1* mutation occurs at high frequency in myelodysplastic syndrome (MDS) and acute myeloid leukemia (AML), the only isogenic *SF3B1* mutant myeloid cell line is in K-562, a chronic myelogenous leukemia (CML) cell line^4^. Conversely, the pre-B ALL-derived Nalm-6 cell line^4^ is the most commonly used B cell line model for *SF3B1* mutation despite *SF3B1* being recurrently mutated in chronic lymphocytic leukemia (CLL)^16^, a mature B malignancy.

**Figure 1:**
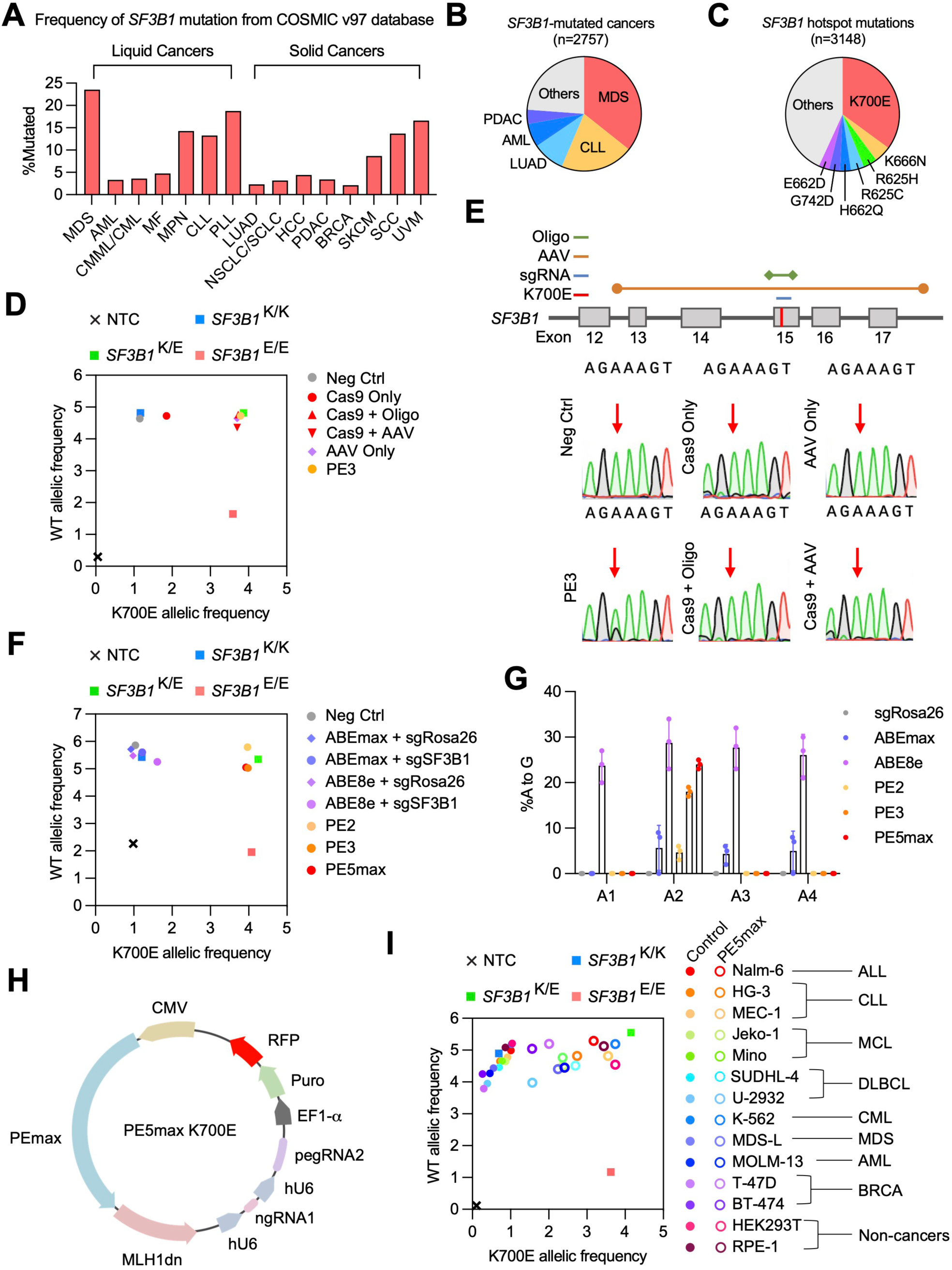
Prime editing can engineer the *SF3B1* mutation efficiently and across spectrum of cell lines and lineages. A) Frequency of *SF3B1* mutation across different cancers and malignancies based on COSMIC v97. The cancers and malignancies shown are: Myelodysplastic syndrome (MDS); Acute myeloid leukemia (AML); Chronic myelomonocytic/myelogenous leukemia (CMML/CML); Myelofibrosis (MF); Myeloproliferative neoplasms (MPN); Chronic lymphocytic leukemia (CLL); Prolymphocytic leukemias (PLL); Lung adenocarcinoma (LUAD); Non-small cell carcinoma (NSCLC/SCLC); Hepatocellular carcinoma (HCC); Pancreatic ductal adenocarcinoma (PDAC); Breast cancer (BRCA); Skin cutaneous melanoma (SKCM); Squamous cell carcinoma (SCC); Uveal melanoma (UVM). All data were derived from the COSMIC database. B) All cancers and malignancies containing *SF3B1* mutation represented by proportion of total number of mutated cases from COSMIC v97. C) The most common *SF3B1* mutation with associated amino acid changes and frequency from COSMIC v97. D) Allelic discrimination plot comparing K700E editing for PE3 versus Cas9 and AAV-homology directed repair in HEK293T based on rhAMP SNP assay. E) (top) Scheme showing the editing strategies for inserting the K700E mutation using Cas9 and AAV-HDR and (bottom) Sanger sequencing results from HEK293T transfected with indicated constructs. Red arrow indicates the nucleotide targeted for editing (AAA>GAA). F) Allelic discrimination plot comparing K700E editing for different prime editing systems versus base editors in HEK293T based on rhAMP SNP assay. G) Editing efficiency of experiment performed in F) measured through Sanger sequencing in biological triplicates. H) The design of the PE5max K700E all-in-one construct. A cassette with pegRNA and ngRNA under hU6 promoters along with a downstream ORF containing puromycin selection marker and TagRFP is cloned into a backbone containing PEmax and MLH1dn. I) Allelic discrimination plot showing PE5max editing of K700E in a variety of cell lines based on rhAMP SNP assay. Cell lines belonging to different cancers include: Acute lymphoblastic leukemia (ALL), Chronic lymphocytic leukemia (CLL); Mantle cell lymphoma (MCL), Diffuse large B cell lymphoma (DLBCL); Chronic myelogenous leukemia (CML); Myelodysplastic syndrome (MDS); Acute myeloid leukemia (AML); Breast cancer (BRCA).

The lack of proper isogenic models is attributable to several factors, with one key culprit being the limitations of current site-specific mutagenesis methods. Most *SF3B1* mutant cell lines are engineered via CRISPR-Cas9 or AAV technologies^3–5,17,18^. Both approaches utilize homology-directed repair (HDR) to insert the K700E mutation into *SF3B1*, which relies on cell’s intrinsic DNA repair capacity for efficient editing and underlies the limited availability of *SF3B1* mutant cell lines. Because *SF3B1* is essential to cell survival, engineering approaches that use double stranded DNA breaks (DSBs) as prerequisite to editing are severely penalized; This is further confounded by most cancer cell lines often exhibiting defective DNA repairs. Furthermore, efficient HDR is heavily dependent on *TP53* status that is frequently impacted across different tissues and cancers cell lines^19,20^. These factors all contribute to the limited availability of disease-relevant *in vitro* models.

Novel and emergent CRISPR-based technologies such as base editing and prime editing edit the genome through non-DSB approaches and therefore, are less cytotoxic and more efficient than Cas9 HDR^21^. Base editing pairs a nickase Cas9 (nCas9) with either a cytosine or adenosine deaminase to effectuate CRISPR-directed DNA editing in the genome^22^. In this capacity, base editing has been proven effective in targeting splicing genes for mutagenesis^23^. However, base editing suffers from poor mutagenic accuracy, rendering certain types of allelic edits impossible^24,25^. In contrast, prime editing effectuates genome-editing through CRISPR-directed reverse transcription^26,27^; A prime editing guide RNA (pegRNA) initiates editing by directing the nCas9 to nick single-stranded DNA that is then primed by the pegRNA for reverse transcription to incorporate edits into the genome. As a result, prime editing is more accurate than base editing while remaining less cytotoxic and more efficient than Cas9 and AAV-based approaches^26^.

In this study, we show that prime editing can efficiently engineer the oncogenic *SF3B1* K700E hotspot mutation into cell lines, outperforming contemporary Cas9, AAV, and base editing approaches. We further show that prime editing can be coupled with synthetic intron splicing reporters to facilitate cell line engineering that we used to create *SF3B1* mutant CLL cell lines. Finally, we demonstrate that these CLL *SF3B1* mutant cell lines accurately model for *SF3B1* mutated CLL, highlighting prime editing as a powerful tool for future pathogenic studies.

## Methods

### Molecular cloning

All primers used for cloning are listed in **Supplementary Table 2**. To generate a *SF3B1* K700E positive control for rhAMP SNP assays, a FragmentGene (GeneWiz) containing the K700E mutation (AAA>GAA) plus 350bp of upstream and downstream sequences was amplified using Q5 polymerase (New England Biolabs) and cloned into pUC19 (Invitrogen) between HindIII and BamHI (New England Biolabs). The pUC19-*SF3B1*-K700E plasmid is also used as a double-stranded DNA (dsDNA) template for Cas9 homology-directed repair (HDR). All oligos for sgRNA cloning were synthesized by Integrated DNA Technologies. All sgRNA and pegRNA sequences are listed in **Supplementary Table 3**. Cas9 HDR sgRNA sequence was designed using HDR Donor Designer (Horizon Discovery) and ligated into pSpCas9(BB)-2A-GFP (Addgene plasmid 48138) between BbsI (New England Biolabs) cut sites to produce the *SF3B1* sgRNA Cas9-GFP construct. Single-stranded DNA (ssDNA) oligo with *SF3B1* K700E mutation to serve as repair template for Cas9-HDR was generated through the Alt-R CRISPR service (Integrated DNA Technologies). Base editing sgRNA directed against the *SF3B1* K700 locus was designed using BE-Designer^28^ and cloned into pLKO.5 Puro-2A-GFP between BsmBI (New England Biolabs) cut sites. pegRNAs and ngRNAs were designed using PegIT^29^. All pegRNAs were ligated into the pU6-pegRNA-gg-acceptor vector (Addgene plasmid 132777) between BsaI (New England Biolabs) cut sites as described^26^. All ngRNAs were cloned into pLKO.5 Puro-2A-RFP between BsmBI (New England Biolabs) cut sites, resulting in the ngRNA1 or ngRNA2 pLKO.5 Puro-2A-RFP backbone. To construct the pCMV-PE5max-*SF3B1*-K700E vector (**Fig. 1h**), the hU6-pegRNA2-polyT cassette was amplified from the pU6-pegRNA-gg-acceptor backbone and ligated into ngRNA1 pLKO.5 Puro-2A-RFP between EcoRI and XhoI (New England Biolabs) to create pLKO.5-K700E-ng1+pg2-Puro-2A-RFP. The hU6-ngRNA1-hU6-pegRNA2-EF1-α-Puro-2A-RFP-WPRE cassette was amplified from this backbone and ligated into pCMV-PEmax-P2A-hMLH1dn (Addgene plasmid 174828) between SgrDI (Thermo Fisher Scientific) cut sites. To generate the pLenti-*SF3B1*-K700E-reporter, the *MTERF2* synthetic intron (synMTERFD3i1-150)^30^ was used to split EGFP between residues Q95 and E96 to create the EGFP synthetic intron reporter (EGFPi). An EF1-α promoter was used to drive the biscistronic expression of a blasticidin resistance marker (BSD) and EGFPi separated by a 2A autocleavable signal (**Fig. 2a**). This cassette was placed in an anti-sense orientation between two LoxP sequences and a WPRE signal and cloned into the pTwist+lenti+SFFV backbone (Twist Biosciences) between BamHI and XhoI (New England Biolabs) cut sites. The SFFV promoter is removed using Q5 site-directed mutagenesis (New England Biolabs). All plasmid constructs were verified using Primordium whole plasmid sequencing (Primordium Labs).

**Figure 2:**
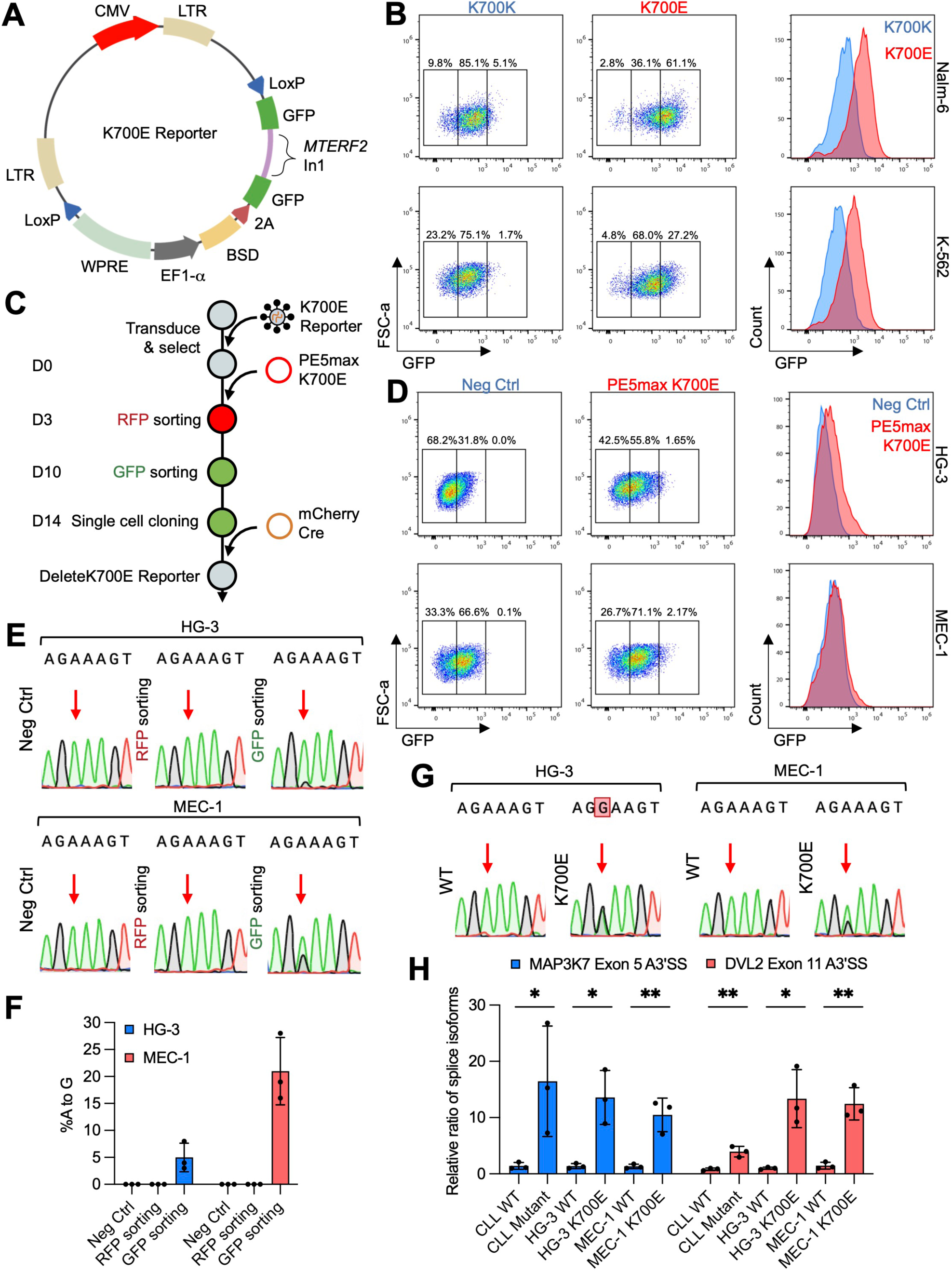
*SF3B1* mutation-responsive reporter allows for facile isolation of *SF3B1* mutant clones. A) The design of the K700E reporter. GFP is bisected by synMTERFD3i1-150^30^ and co-expressed with a blasticidin selection marker. The entire cassette is flanked by loxP signals to enable Cre recombinase-mediated deletion of reporter post-editing. B) Flow cytometry plots (left) and histograms (right) of Nalm-6 and K562 *SF3B1* K700K and K700E cell lines stably transduced with the K700E reporter. C) Workflow for engineering cell lines with the K700E mutation. D) Flow cytometry plots (left) and histograms (right) of HG-3 and MEC-1 control cells or cells electroporated and sorted for PE5max K700E. E) Sanger sequencing results for HG-3 and MEC-1 control cells, cells sorted for RFP (PE5max K700E), and cells sorted for GFP (K700E reporter). F) K700E frequency measured through Sanger sequencing for each sample in biological triplicates. G) Sanger sequencing results for isogenic HG-3 and MEC-1 *SF3B1* WT and K700E clones. H) Splice variant qPCR analysis of *MAP3K7* and *DVL2* alternative splicing in CLL, HG-3, and MEC-1 with and without *SF3B1* mutation in biological triplicates. Student unpaired t-test was used (*P≤0.05, **P≤0.01).

### Human Samples

Blood samples were withdrawn with informed consent from patients enrolled in clinical protocol as approved by the Human Subjects Protection Committee of City of Hope. To isolate B cells, peripheral blood mononuclear cells were enriched through density gradient centrifugation followed by immuno-magnetic negative selection using Pan B cell isolation kit (MiltenyBiotec). Samples are then cryo-preserved until needed for study. *SF3B1* mutation was verified in patient samples through a targeted sequencing panel.

### Cell lines and culture

HEK293T, RPE-1, and BT-474 cells were cultured in DMEM (Gibco) with 10% FBS (Omega Scientific) and 100ug/ml penicillin-streptomycin (Gibco). T-47D and Nalm-6 were cultured in RPMI (Gibco) with 10% FBS (Omega Scientific) and 100ug/ml penicillin-streptomycin (Gibco). HG-3, MEC-1, Jeko-1, Mino, SUDHL-4, U-2932, MDS-L, and MOLM-13 were cultured in RPMI (Gibco) with 20% FBS (Omega Scientific), 100ug/ml penicillin-streptomycin (Gibco), and 2mM supplemented glutamine (Gibco). MDS-L was co-cultured with 30ng/ml human IL-3 (Prepotech). K-562 was cultured in IMDM (Gibco) with 10% FBS (Omega Scientific) and 100ug/ml penicillin-streptomycin (Gibco). BT-474 and T-47D were generously gifted by Dr. Chun-Wei Chen (City of Hope). RPE-1 was generously gifted by Dr. Jeremy Stark (City of Hope). MDS-L was generously gifted by Dr. Ling Li (City of Hope). MOLM-13 was generously gifted by Dr. Jianjun Chen (City of Hope). Nalm-6 and K-562 cell lines endogenously expressing *SF3B1* K700K or K700E were purchased from Horizon Discovery. HEK293T with endogenous expression of *SF3B1* K700E was generated through prime editing. HEK293T, HG-3, and MEC-1 WT isogenic controls are cells that do not have the K700E mutation but arose through single cell cloning alongside the K700E clones. Adherent cells were maintained on 6-well plates while suspension cells were grown on 12-well plates. All cell lines were regularly tested and confirmed to be negative for mycoplasma.

### AAV and lentiviral packaging

AAV packaging and purification was done as described^31^. The pAAV-SEPT-*SF3B1*-K700E donor plasmid and pRC1 and pHelper packaging plasmids were generously donated by Dr. Brian William Dalton (Johns Hopkins Medicine)^3^ and were transfected into HEK293T at 80% confluency in 10cm plate in a 1:1:1 ratio of 10ug each. 3 days after transfection, the AAV packaging and purification were done as described^31^. Briefly, AAV-*SF3B1*-K700E virions were released from HEK293T using the freeze-thaw method and precipitated overnight at 4°C using PEG 8000 (Fisher Scientific). Precipitates were pelleted via centrifugation at 2800 g for 15min at 4°C, resuspended in 2ml PBS, and then incubated with 2ul of DNase I (New England Biolabs), 2ul RNase A (Invitrogen), and 3.5ul of 1M MgCl_2_ (Invitrogen) for 20min at 37°C. Viral supernatant was then mixed 1:1 with chloroform and incubated for 30 min at room temperature. After centrifuging at 3000 g for 15min at room temperature, the top aqueous layer was isolated and passed through an Amicron Ultra-0.5 filter (MilliporeSigma). PBS was passed through the column twice to wash the concentrated virus and to remove chloroform. The final concentrated viral was eluted from the column by inverting the column into a 1.5ml tube and centrifuging at 1000 g for 1min. For AAV HDR experiments, HEK293T were pre-seeded onto 12-well plates and grown to 80% confluency before transduction by addition of the concentrated virus to the media. For combination with Cas9 editing, HEK293T was transduced 24 hours before transfecting in *SF3B1* sgRNA Cas9-GFP. To engineer cell lines harboring the pLenti-*SF3B1*-K700E-reporter, cells were transduced via spin-infection. To package lentivirus, 3 ug of viral plasmid donor, 0.4 ug of VSV-G, and 1.5 ug psPAX2 were mixed with PEI-MAX 40K (Polyscience) and transfected into seeded HEK293T in 6-well format. Lentiviral supernatant was collected at 72hrs and 96hrs after transfection, pooled, and filtered through a 0.45um PES syringe filter (Bioland Scientific) before being concentrated with ultracentrifugation at 88,000 g for 2hrs at 4°C. To transduce suspension cells, 0.5 million cells were seeded into a 48-well format with 8ug/ml polybrene and concentrated lentivirus and spin-infected at 1000 g for 1.5 hours at 37°C. The media was changed 3 hours after the spin-transduction. Transduced cells were then selected with 10ug/ml blasticidin (InvivoGen) for 5 days.

### Transfection and flow cytometry

For Cas9 and AAV HDR, base editing, and prime editing experiments in HEK293T cells, 50,000 cells were seeded onto 12-well plates, grown to 80% confluency, and transfected using PEI MAX 40K (Polysciences). For Cas9 and AAV HDR experiments, 1ug of *SF3B1* sgRNA Cas9-GFP was transfected alone if cells were pre-transduced with AAV-*SF3B1*-K700E or with either 0.1nmole of the *SF3B1* Alt-R HDR template. For base editing experiments, 1ug of either *SF3B1* sgRNA or *Rosa26* sgRNA in pLKO.5 Puro-2A-GFP backbone was co-transfected with either 1ug of NG-ABE8e (Addgene plasmid 13849) or NG-ABEmax (Addgene plasmid 124163). For prime editing experiments, 0.5ug of pegRNAs were co-transfected alone or with 0.5ug ngRNA with 1ug of either pCMV-PE2-P2A-GFP (Addgene plasmid 132776) or pCMV-PEmax-P2A-MLH1dn (Addgene plasmid 174828). HEK293T, RPE-1, T-47D, and BT-474 were transfected using PEI MAX 40K (Polysciences). For prime editing with the pCMV-PE5max-*SF3B1*-K700E all-in-one construct, 3ug of the plasmid was transfected into HEK293T, RPE-1, T-47D, and BT-474 on 6-well plates at 80% confluency. Nalm-6, HG-3, MEC-1, Jeko-1, Mino, SUDHL-4, U-2932, MDS-L, MOLM-13, and K-562 were electroporated using the Celetrix LE+ system (Celetrix). For electroporation, ∼30-40 million cells were resuspended in 200ul volume of electroporation buffer and 30ug of plasmids and electroporated at 1100V, 30ms, 1x pulse. Cells were allowed to recover for 3-4 days before being taken for genomic DNA extraction. For testing Cas9 HDR in HG-3 cells, Alt-R Cas9v3 and *SF3B1* sgRNA RNP (Integrated DNA Technologies) were assembled into RNPs following manufacturer’s instruction. Then, 5 million cells were electroporated with either 0.125nmole of assembled RNP and 0.1nmole of the *SF3B1* Alt-R HDR template or 10ug of *SF3B1* sgRNA Cas9-GFP plasmid and 30ug of the pUC19-*SF3B1*-K700E plasmid as a dsDNA repair template. HG-3 cells were electroporated using either Neon (Invitrogen) in the case of RNP or the Celetrix LE+ system (Celetrix) in the case of plasmid. For the former, the following settings were used to get >99% electroporation efficiency: 1600V, 10ms, 3x pulses. For the latter, cells were electroporated using 1100V, 30ms, 1x pulse followed by cell sorting using an Aria Fusion (BD Biosciences). To engineer *SF3B1* K700E cell lines via prime editing, ∼30-40 million cells harboring the pLenti-*SF3B1*-K700E-reporter were electroporated with 20ug of the pCMV-PE5max-*SF3B1*-K700E and expanded over 3 days before being sorted for RFP for which at least 200,000 events were collected. Sorted cells were re-seeded into pre-warmed media in a 96-well format and expanded over 7 days before being sorted for GFP, collecting at least 200,000 events. Sorted cells were then re-seeded into pre-warmed media in a 96-well format and expanded for 4 days before being taken for single cell cloning. Some cells were collected at day 7 after RFP sorting and day 4 after GFP sorting for genomic DNA isolation to check for presence of the K700E mutation. To remove pLenti-*SF3B1*-K700E-reporter, 20ug of pCMV-mCherry-Cre (Addgene plasmid 27546) was electroporated into ∼30-40 million cells using the setting 1100V, 30ms, and 1x pulse and sorted for RFP. Electroporated cells were recovered, expanded, and sorted again for GFP negative cells followed by single cell cloning to isolate pure clones. All flow cytometries were performed on a Fortessa X20 (BD Biosciences), and flow data were analyzed using FlowJo version 10.8.2 (BD Biosciences).

### Genetic analysis

Genomic DNA were extracted from cells using the NucleoSpin Tissue kit (Macherey-Nagel) and stored in TE buffer. In experiments involving high-throughput screening of single cell clones, genomic DNA was released from cells by thermolysis. Briefly, cells were spun down in 96-well PCR plates (Bioland Scientific), resuspended in 20ul water, and heated at 100°C for 10min in a thermocycler (Applied Biosystems). After pelleting cellular debris, the supernatant was used for genotyping. To check for the K700E mutation via Sanger sequencing, the *SF3B1* K700 locus was amplified by PCR on genomic DNA using DreamTaq (Invitrogen) with the primer pairs: forward, 5’ CCTGTGTTTGGTTTTGTAGGTC 3’ and reverse, 5’ GGTGGATTTACCTTTCCTCT 3’. In the experiments where presence of residual Alt-R HDR oligos or AAV-*SF3B1*-K700E DNA can result in false positives, a different set of PCR primers that will only amplify genomic DNA were used: forward, 5’ GTGTAACTTAGGTAATGTTGGGGC 3’ and reverse, 5’ GAAGAGAAAAGTGACCAAACATCG 3’. PCR products were purified using the NucleoSpin Gel and PCR Clean-up kit (Macherey-Nagel) and sequenced using Sanger sequencing (Eton Biosciences) using the reverse primer for PCR done with the first primer pairs and the forward primer for PCR done with the second primer pairs. The *SF3B1* genetic sequences was downloaded from the GRCh38 reference using IGV (Broad Institute) and was used to align the Sanger sequencing trace files with SnapGene (Dotmatics). Editing efficiency was calculated using ICE^32^ (Synthego) and EditR^33^ (http://baseeditr.com) on biological triplicates. Deletion of the pLenti-*SF3B1*-K700E-reporter via Cre recombinase was confirmed with PCR using two pairs of primers: PCR1 - forward, 5’ CTTGCTCACCATCGGTCCAG 3’ and reverse, 5’ ATGGCCAAGCCTTTGTCTCAAG 3’; PCR2 - forward, 5’ ACTGCTGATCGAGTGTAGCC 3’ and reverse, 5’ TCATTGGTCTTAAAGGTACCGAGC 3’. PCR1 and PCR2 were performed at the annealing temperatures of 61°C and 58°C, respectively, with an extension time of 30 seconds at 72°C. Cre recombinase-mediated deletion of the K700E reporter was assessed on 2% agarose gel in 1xTAE buffer with a 1kB Plus Ladder (Invitrogen). RhAMP SNP primers (Integrated DNA Technologies) were designed and used to genotype cells. For clean genomic DNA isolated using kit, ∼100-200ng was used as inputs for the PCR reaction. For crude genomic DNA released through thermolysis, 4.2ul of the supernatant was used. For experiments where residual Alt-R HDR oligos or AAV-*SF3B1*-K700E DNA may result in false positives, the *SF3B1* K700 locus was first amplified with the *SF3B1* AAV primer set to generate purified PCR products for rhAMP SNP assay. For rhAMP SNP assay on PCR products, 10pg of PCR products were used. For rhAMP SNP assay on gDNA, 100ng of gDNA was used. 1ng of pUC19-*SF3B1*-K700E and 100ng of genomic DNA from K562 *SF3B1* K700K and K700E were used as controls for homozygous and heterozygous mutants and WT, respectively. RhAMP SNP primers were designed by IDT DNA (assay name CD.GT.WVSM3699.1). RhAMP SNP assay was carried out per manufacturer’s protocol on a QuantStudio 12K Flex (Applied Biosystems) in a 384-well format (USA Scientific). Allelic discrimination was plotted using Graphpad Prism (Dotmatics). All primers used in PCR and genotyping are listed in **Supplementary Table 4**.

### Splice variant qPCR

Total RNA was extracted using Trizol (Ambion) while cDNA was generated using the High-Capacity cDNA Reverse Transcription Kit (Applied Biosystems) per manufacturer’s protocol. 5ng of total cDNA was used as inputs for qPCR on the QuantStudio 12K Flex (Applied Biosystems). To calculate relative ratio of splice variants, The Ct values for the normal and alternatively spliced isoforms were inputted into the following formula: 2^ΔΔCt(Normal)-ΔΔCt(Aberrant)^. The results were graphed using GraphPad Prism (Dotmatics). For *MTERF2* intron 1 splicing analysis, 5ng of total cDNA was used as inputs for PCR reactions using three primers: *MTERF1* Ex1, 5’ ACTCCCTGTGCCTTGCTTG 3’; *MTERF1* In1, 5’ CATTTTGGGCATGGAATCTG 3’; *MTERF1* Ex2-3 R, 5’ ATGGACTCATTCTATCTTACAGTCTCTCC 3’. All PCR products were resolved on 2% agarose gel in 1xTAE buffer with a 1kB Plus Ladder (Invitrogen). Primers to detect splice variants are listed in **Supplementary Table 5**.

### RNA-seq and RNAseqCNV analysis

PolyA mRNA was enriched from total RNA and constructed into a library for RNA-seq using the Stranded Total RNA Prep with Ribo-Zero Plus Kit (Illumina). cDNA libraries were sequenced using paired-end 150bp on the Illumina NovaSeq 6000 platform at 50 million reads per samples. Reads were aligned to the GRCh38 reference genome using STAR. Splicing analysis was performed using rMATS^34^. RNA-seq was carried on HG-3 and MEC-1 clones that have underwent Cre recombination to remove the K700E reporter and isolated through single-cell cloning. RNA-seq data for Nalm-6 and K562 *SF3B1* K700K and K700E cell lines were downloaded from NCBI GEO under accession GSE72790. For primary CLL patient samples, 23 *SF3B1* WT and 13 *SF3B1* mutant samples were used for RNA-seq analysis and were previously deposited^35^. RNA-seq read coverage plots were generated by plotting RNA-seq data using IGV (Broad Institute). For splice variant analysis in primary patient samples, FDR ≥0.05, abs(ΔPSI) ≥0.1, and total_reads ≥3 were used as statistical cutoffs. For each cell lines, different reads cutoffs were used to normalize for differential total sample reads between cell lines: Nalm-6, total_reads ≥3; HG-3, total_reads ≥10; MEC-1, total_reads ≥15. rMATS output for CLL, Nalm-6, HG-3, and MEC-1 are shown in **Supplementary Tables 6-9**. RNA-seq data was used to analyze for CNVs in HG-3, MEC-1, and Nalm-6 *SF3B1* WT and K700E cell lines using RNAseqCNV^36^.

### Whole genome sequencing and copy number variation detection

Genomic DNA were isolated from cell lines using the NucleoSpin Tissue extraction kit (Macherey-Nagel). Genomic DNA amount and quality were assessed using a NanoDrop (Thermo Fisher) and taken for whole genome sequencing with Nebula Genomics at 3-18x coverage. The whole-genome sequencing fastq files were aligned to the GRCh38 reference genome using BWA-MEM^37^ to generate BAM files. Subsequently, each BAM file underwent sorting and indexing through SAMtools^38^. Duplicate reads were identified and marked using the MarkDuplicates function in Picard (https://broadinstitute.github.io/picard/). SAMtools^38^ were then employed to generate an index for each BAM file with marked duplicate reads. The entire genome was segmented into windows with a width of 10,000 bp. The read depth within each window was extracted using mosdepth^39^, producing a BED file that captures window start, end, and depth information. Utilizing BED files, copy number variation detection was performed using the R package cn.mops^40^, comparing the tumor sample against the wild-type sample through referencecn.mops and calcIntegerCopyNumbers functions. The outcome is a copy number variation table detailing the start, end, and the observed copy number variations. Whole-genome sequencing data for primary CLL patients were downloaded from CLLmap^41^.

### Cancer distribution and frequency of *SF3B1* mutation

The cancer distribution and frequency of *SF3B1* mutation were downloaded from COSMIC v97 (cancer.sanger.ac.uk) on January 28^th^, 2023. COSMIC data was filtered for point mutations on *SF3B1* (COSG68561) and for cancers with at least >2% of samples with *SF3B1* mutation. For **Fig. 1a**, the percentage of samples with *SF3B1* mutation for each cancer was calculated and plotted using GraphPad Prism (Dotmatics). For **Fig. 1b** and **Fig1.c**, the total numbers of samples with *SF3B1* mutation for each cancer types and samples with a specific *SF3B1* mutation were plotted as pie charts using GraphPad Prism (Dotmatics). Venn diagrams were created using the Multiple List Comparator from MolBioTools (www.molbiotools.com/listcompare).

### Statistical Methods

All statistical tests were calculated using Graphpad Prism (Dotmatics). Error bars are shown to represent the mean of three independent replicates. For **Fig. 2h** and **Fig. 3g**, unpaired t-test was performed to calculate P-values between biological triplicates of *SF3B1* WT or mutant from primary CLL patient samples and cell lines. For supplementary Fig. 8a-d, the one sample Wilcoxon signed rank test was performed for each chromosomal location using log_2_FC=0 as the hypothetical value. For **Fig. 4d**, a Mann-Whitney test was performed between *SF3B1* WT and mutant primary CLL patient samples for CNV calls from WGS. For **Fig. 4e,f**, a chi-squared test was performed with -log_10_(0.05) chosen as the statistical cutoff value.

**Figure 3:**
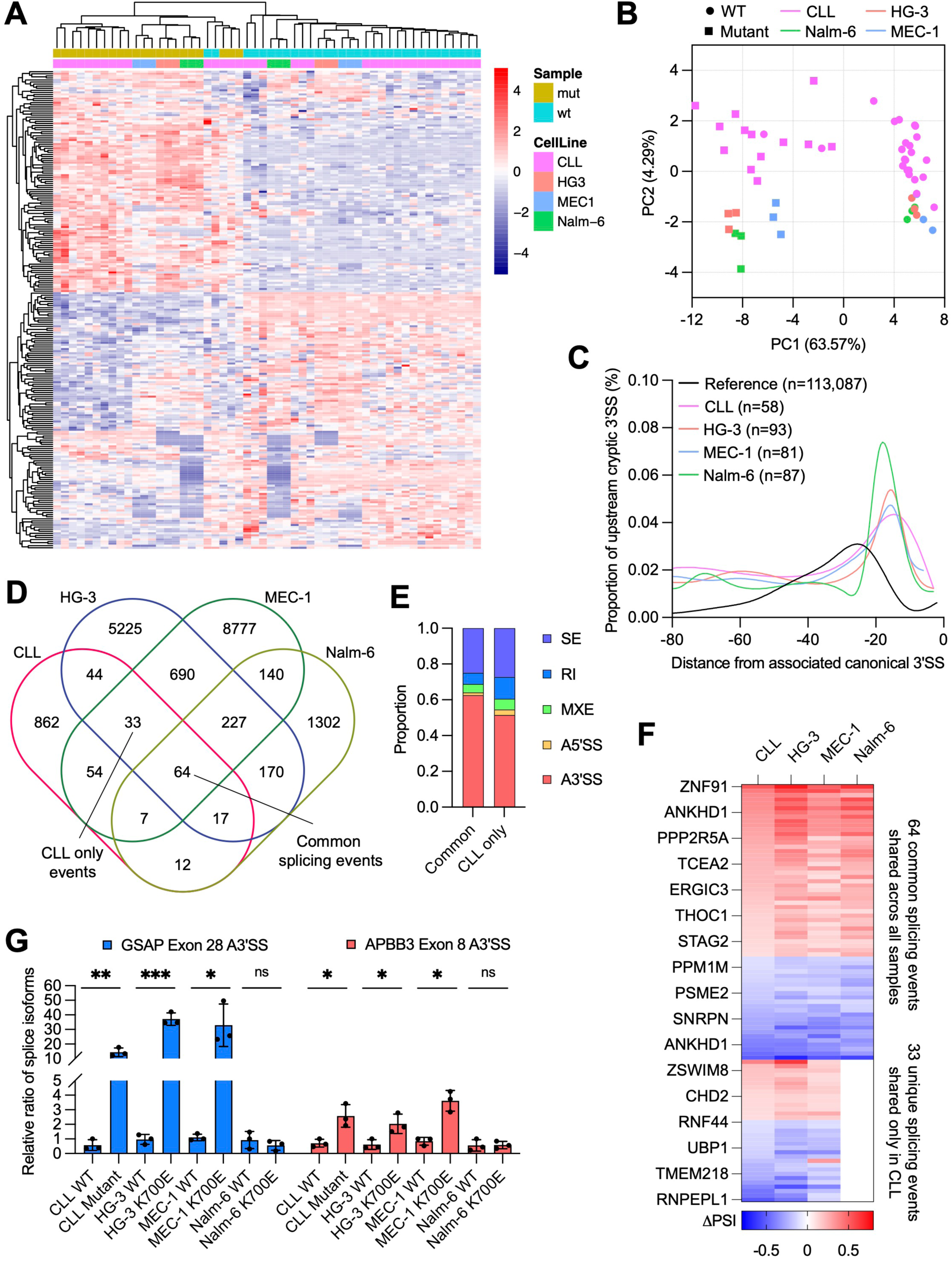
CLL *SF3B1* mutant cell lines capitulates the altered splicing profile of primary CLL. A) Unsupervised clustering analysis and B) PCA plot based on alternative splicing events of RNA sequencing data derived from primary CLL patient samples (n=36) and cell lines with and without *SF3B1* mutation. C) Density plot showing the position and frequency of cryptic 3’SS for *SF3B1*-mutated primary CLL samples and cell lines versus average distance to the first AG from RefSeq reference of all canonical 3’SS. D) Overlap analysis of significant splicing events associated with *SF3B1* mutation (PSI>0.1 or <-0.1 with P value<0.05) among CLL, HG-3, MEC-1, and Nalm-6 cells. Events common to all samples or specific only to CLL and CLL cell lines are indicated. E) Stacked bar plot representation of the type of alternative splicing events found in common and CLL only splice variants. F) Heatmap showing the splicing profiles of 64 common and 33 CLL only splice variants. G) Splice variant qPCR analysis of *GSAP* and *APBB3* alternative splicing in CLL, HG-3, MEC-1 and Nalm-6 with and without *SF3B1* mutation in biological triplicates. Student unpaired t-test was used (*P≤0.05, **P≤0.01).

**Figure 4:**
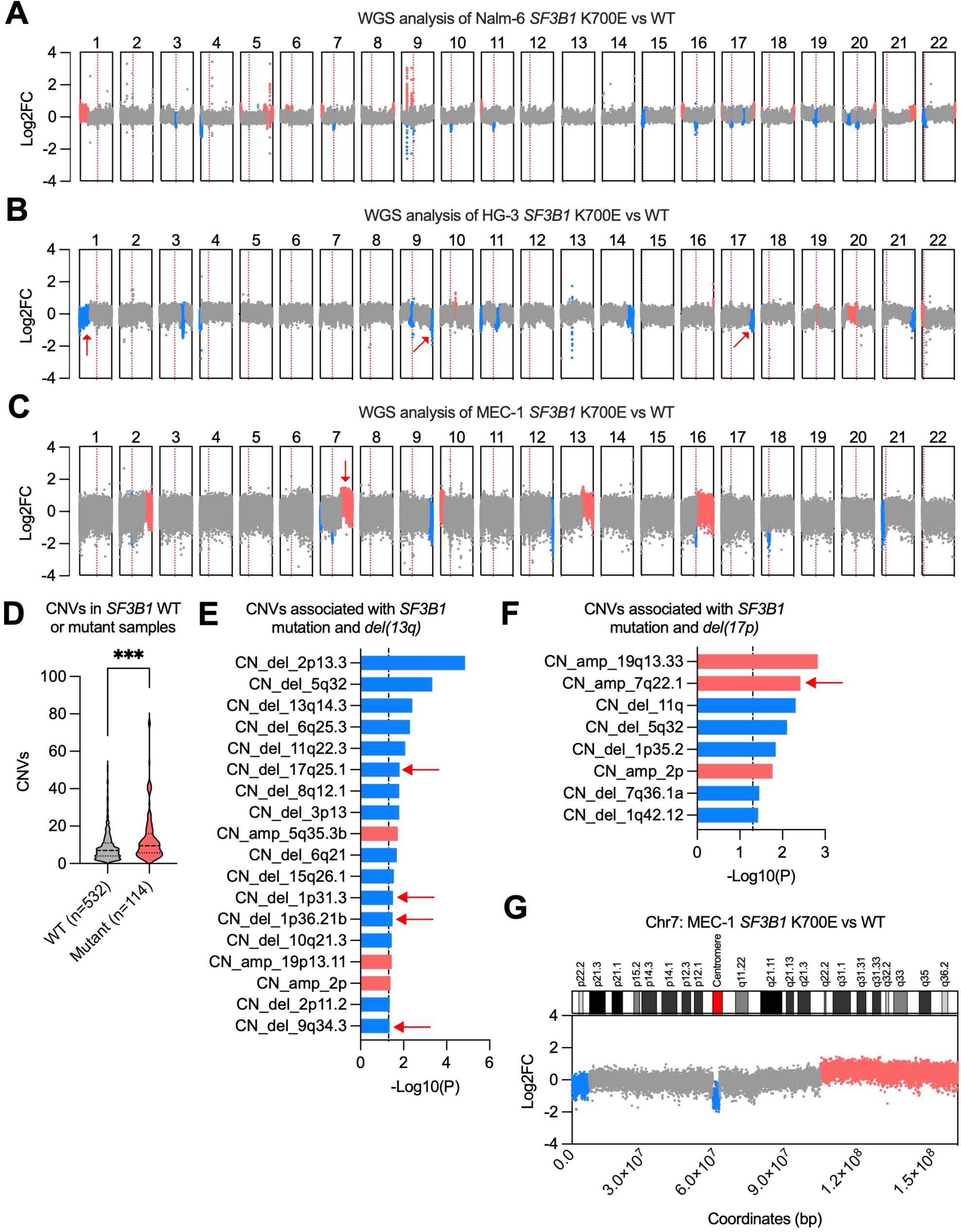
*SF3B1* mutant cells exhibit significant CNV changes that are consistent with primary CLL. CNVs analysis comparing *SF3B1* mutant samples versus isogenic WT based on whole-genome sequencing data for A) Nalm-6, B) HG-3, and C) MEC-1. Significant CNV events are indicated by blue for deletions and red for amplifications. Centromeric region demarcating the p and q chromosomal arms is indicated by a red dotted line. Red arrows indicate CNV events shared with primary patient CLL. D) Violin plot of the number of CNV events found in *del*(17p) and *del*(13q) CLL patients that are either *SF3B1* WT or mutant based on CLLmap data (https://cllmap.org/). A Mann-Whitney test was used (***P≤0.001). CNVs associated with either E) *SF3B1* mutation and *del*(13p) and F) *SF3B1* mutation and *del(17p)* co-occurrences. Red arrow indicates chromosomal amplification shared between primary CLL samples and cell lines. A chi-squared test was performed with - log_10_(0.05) as the statistical cutoff. G) (top) Cytogenetic loci on the p and q arm and (bottom) WGS analysis for chromosome 7.

## Data Availability

Constructs used in this study will be made available on Addgene (www.addgene.org/Lili_Wang/). All RNA-seq results for the HG-3 and MEC-1 *SF3B1* WT and K700E cell lines have been deposited in the Gene Expression Omnibus (GEO) database with accession number GSE231387.

## Acknowledgements

This work was supported by fundings from the NIH/NCI R01CA240910 and R01CA21623 (to L.W.) and F31CA261110 (to M.F) with additional fellowship funding support from the Held Foundation and H.N. & Frances Berger Foundation (to M.F). The authors would like to acknowledge the helpful advice from Dr. Chih-Hong Lou (City of Hope, Gene Editing and Viral Vector Core) on proper AAV packaging and reagents. Some figures were created using BioRender.

## Author contributions

M.F., R.L., and L.W. designed the study. M.F. performed most of the experiments. K.H. performed some early Cas9 experiments. Q.J., L.Y., X.W., and Z.J. performed the computational analysis. M.F., R.L., and L.W. wrote the manuscript with inputs from all the authors.

## Results

### Prime editing can efficiently engineer the *SF3B1* K700E hotspot mutation

The initial prime editing system, also known as PE2, pairs a prime editing enzyme composed of an nCas9 and an reverse transcriptase with a pegRNA^26^. PE3 improves upon PE2 by using an additional nicking guide RNA (ngRNA) to introduce a secondary nick following the initial edit to bias DNA repair towards incorporating the edit into the genome^26^. We assessed the efficiency of PE2 and PE3 in engineering the K700E hotspot mutation by introducing A>G transition on the first letter in the AAA three letter lysine codon (Supplementary Fig. 1a). We designed pegRNAs and ngRNAs directed against the K700 locus and measured the editing efficiency in HEK293T cells using rhAMP SNP assay and Sanger sequencing (Supplementary Fig. 1b-d). Using these methods, we observed the highest editing efficiency with the PE3 system using the pegRNA2 and ngRNA1 combination (Supplementary Fig. 2a,b). We further benchmarked PE3 against conventional Cas9 and AAV HDR approaches where we observed PE3 introducing the K700E mutation at a higher frequency and lower indels (**Fig. 1d,e**). Encouraged by these results, we next compared PE2 and PE3 to one of the most recent iterations of the prime editing technology, PE5max. Recent works have shown that mismatch repair (MMR) is antagonistic to prime editing^42^; Temporary inhibition of MMR can favor incorporation of edits without causing microsatellite instability^42^. The PE5max improves prime editing by co-expressing an evolved prime editor (PEmax) with a dominant negative isoform of MLH1 (MLH1dn) to temporarily inhibit MMR (Supplementary Fig. 1c). In our tests, PE5max outperformed both PE2 and PE3 in installing K700E mutation on *SF3B1*, achieving up to 25% efficiency in HEK293T cells (**Fig. 1f,g** and Supplementary Fig. 2c,d). Given that *SF3B1* is tr-allelic in HEK293T and that homozygous expression of mutant *SF3B1* is generally lethal, this translates into an overall editing efficiency of 75%.

We next compared prime editing to base editing, an orthogonal gene editing technology. Unlike prime editor, base editors use adenosine deaminases to directly convert A>G on DNA in an sgRNA-dependent manner^43^. We assessed two base editors, ABEmax and ABE8e, and compared their editing efficiency versus PE5max for *SF3B1* K700E editing in HEK293T cells. At the first adenosine position, PE5max attained A>G editing efficacy comparable to the two base editors, ABEmax and ABE8e (**Fig. 1f,g** and Supplementary Fig. 2c,d). However, the base editors also induced high levels of bystander edits to adjacent adenosines resulting in unintended missense mutations. Because the K700 and adjacent codons are adenosine-enriched, this locus is largely uneditable by base editors due the poor mutagenic accuracy, rendering prime editing as the only system that can engineer the K700E mutation. These results highlight the general specificity of prime editing over base editing and demonstrate PE5max as the most optimal system for *SF3B1* engineering.

While prime editing works remarkably well in HEK293T where MMR is deficient, its effectiveness in other cell lines where MMR is intact can be lacking^27,42^. Considering that inclusion of the MLH1dn MMR inhibitor may allow us to efficiently engineer the K700E mutation in a wider variety of cellular contexts, we first reconstituted the components of PE5max – pegRNA2, ngRNA1, PEmax, and MLH1dn – into an all-in-one construct with puromycin and RFP selection markers (pCMV-PE5max-*SF3B1*-K700E, henceforth referred to as PE5max K700E) (**Fig. 1h**). As a functional test of the PE5max K700E system, we used it to derive a pure isogenic model of *SF3B1* mutation in HEK293T (Supplementary Fig. 2e). Through electroporation or transfection of the PE5max K700E construct, we next assayed for K700E prime editing in a panel of cell lines from diverse cancers such as acute lymphoblastic leukemia (ALL), chronic lymphocytic leukemia (CLL), mantle cell lymphoma (MCL), diffused large B cell lymphoma (DLBCL), chronic myelogenous/myelomonocytic leukemia (CMML), myelodysplastic syndrome (MDS), acute myeloid leukemia (AML), breast cancers (BRCA), as well as non-cancer cell lines HEK293T and RPE-1 (**Fig. 1i**). Approximately 72 hours post electroporation or transfection, we were able to detect the presence of the K700E mutation using rhAMP SNP assay in all cell lines (**Fig. 1i**). This result indicates that *SF3B1* K700E engineering is possible across diverse cellular context via PE5max prime editing.

### Coupling prime editing with a *SF3B1* mutation-responsive reporter enables facile engineering

As a proof-of-concept, we next sought to use this system to address the dearth of *SF3B1* mutant B cell lines. While *SF3B1* mutations occur in both lymphoid and myeloid malignancies, most available *SF3B1* mutant cell lines are in the myeloid lineage with few existing lymphoid models (supplementary table 1). *SF3B1* is recurrently mutated in CLL, a mature lymphoid B cell cancer^11^. In our initial effort to engineer the CLL cell line HG-3 with the *SF3B1* mutation using CRISPR Cas9 technology, we observed no successful cases of editing either using Cas9 ribonucleoprotein (RNP) and single-stranded DNA oligo (ssDNA) (Supplementary Fig. 3a,b) or with Cas9 plasmid and a double-stranded DNA (dsDNA) repair template (Supplementary Fig. 3c,d); In the latter case, we also observed several instances of false positives resulting from random integration of the repair template (Supplementary Fig. 3c,d), thereby demonstrating the pitfalls of Cas9 HDR in engineering the *SF3B1* K700E mutation. We thus turned to prime editing to generate CLL *SF3B1* mutant cell line models using two CLL cell lines with pre-existing CLL-associated cytogenetic lesions: HG-3 (*del13q*, 50% of CLL cases^16^) and MEC-1 (*del17p*, 20% of CLL cases^16^). Following PE5max K700E electroporation, we could detect the K700E mutation using rhAMP SNP assay but not Sanger sequencing (Supplementary Fig. 3e,f). This result indicates that while prime editing can successfully install the K700E mutation in a lymphoid context, the frequency remains too low, thereby increasing the difficulty in the isolation of pure single-cell clones (Supplementary Fig. 3g); Further enrichment steps are needed.

A recent report has shown that the intron 1 of the *MTERF2* gene splices out in a *SF3B1* K700E-dependent manner^30^ (Supplementary Fig. 3h). We verified this phenomenon in a panel of *SF3B1* K700E cell lines (Nalm-6, K-562, and HEK293T) where we observed full *MTERF2* intron 1 removal only in mutant cells while WT cells showed either full intron retention or partial intron splicing at one of the cryptic 3’SS (Supplementary Fig. 3i). This observation led us to hypothesize that this intron could be leveraged as a reporter for the K700E mutation. To test this, we inserted a minimal *MTERF2* intron 1 (synMTERD3i1-150)^30^ between EGFP exons to create a fluorescent minigene reporter (EGFPi) (**Fig. 2a**). The EGFPi cassette is placed into a lentiviral construct next to a blasticidin resistance gene (BSD) under the control of an EF1-α promoter. LoxP sites are included to allow for Cre recombinase-mediated deletion of this cassette. We tested this pLenti-*SF3B1*-K700E-reporter (henceforth referred to simply as K700E reporter) in Nalm-6 and K-562 cells harboring either WT or *SF3B1* K700E mutation (**Fig. 2b**). Presence of the K700E mutation led to an increase in both GFP expression and the proportion of GFP bright cells in both cell lines. Given that this reporter can fluorescently label K700E prime edited cells, we devised an innovative approach termed <u>pr</u>ime <u>e</u>diting <u>c</u>oupled intron-assisted <u>s</u>election (PRECIS) that pairs fluorescent sorting with prime editing (**Fig. 2c** and Supplementary Fig. 3j).

Using the reporter, we observed a noticeable increase in the GFP dim and bright populations in PE5max K700E-electroporated HG-3 and MEC-1 cells when compared to negative control cells (**Fig. 2d** and Supplementary Fig. 4a,b). Following FACS-enrichment of the GFP bright population, we observed detectable levels of A>G edits in HG-3 (∼5%) and MEC-1 (∼22%) through Sanger sequencing (**Fig. 2e,f** and Supplementary Fig. 4c,d). Clonal selection via single-cell cloning paired with high-throughput rhAMP SNP screening yielded HG-3 and MEC-1 clones harboring heterozygous K700E alleles (**Fig. 2g** and Supplementary Fig. 4e,f). For HG-3, we were able to derive pure heterozygous clones while with MEC-1 only aneuploid (tri-allelic) clones were possible. We then removed the K700E reporter using Cre-loxP system through overexpression of Cre recombinase (Supplementary Fig. 5a-d). We further confirmed the phenotypic impact of *SF3B1* mutation by checking for known alternative splicing in these cell lines including full *MTERF2* intron 1 splicing and consistent upregulation of *MAP3K7* and *DVL2* splice variants^7^ (**Fig. 2h** and Supplementary Fig. 6a-b). These results confirmed that we have generated *bona fide* CLL *SF3B1* K700E cell lines.

### CLL *SF3B1* K700E cell lines recapitulate the altered splicing of *SF3B1*-mutated primary CLL

We next sought to benchmark these novel CLL and Nalm-6 *SF3B1* mutant cell lines for their ability to recapitulate primary *SF3B1*-mutated CLL cells. Across primary CLL samples, CLL cell lines, and Nalm-6, presence of *SF3B1* mutation drove a uniform splicing paradigm with all *SF3B1*-mutated samples clustering together (**Fig. 3a,b**). Presence of *SF3B1* mutation promoted selection of cryptic 3’SS proximal to within 15nts of the canonical 3’SS across all samples (**Fig. 3c** and Supplementary Fig. 7a). Mutant *SF3B1* also preferentially affected alternative 3’SS splice site selections as well as intron retention compared to other splicing events (Supplementary Fig. 7b). These results confirmed that the HG-3 and MEC-1 *SF3B1* K700E cell lines shared many similarities with *SF3B1*-mutated primary CLL and Nalm-6 cells. When we compared splice variants associated with the mutation, however, we found both common splice variants shared with both Nalm-6 and primary CLL cells as well as novel splice variants shared only with primary CLL cells (**Fig. 3d**). Splice variants belonging to both common and CLL only groups were predominantly alternative 3’SS splicing events (**Fig. 3e**). Common splice variants showed consistent splicing patterns across all *SF3B1*-mutated samples, confirming a uniformity in *SF3B1* mutation-mediated splicing program while CLL only variants showed splicing between primary CLL and CLL cell lines only, being completely absent in Nalm-6 altogether (**Fig. 3f**). These results suggest lineage-specific alternative splicing for *SF3B1* mutation. To confirm this, we checked for aberrant splicing of two targets that undergo alternative 3’SS splicing, *GSAP* and *APBB3*. Using qPCR, we confirmed upregulated expression of *GSAP* and *APBB3* alternative splice variants only in CLL and CLL cell lines but not Nalm-6 (**Fig. 3g**). These results collectively show that HG-3 and MEC-1 *SF3B1* K700E cell lines not only faithfully recapitulate the altered splicing in *SF3B1*-mutated CLL, but also display CLL-specific splicing events that cannot be modeled in Nalm-6, thereby unlocking unprecedented opportunities for studying the impact of *SF3B1* mutation in CLL leukemogenesis.

### *SF3B1* mutation drives general and CLL-specific CNV events

In CLL, *SF3B1* mutation frequently co-occurs with two common CLL genetic lesions, del*(13q)* and del*(17p)*, that result in ablation of the *BCL2*-targeting *MIR15*/16 locus and *TP53*, respectively^16,41,44^. From the clonal evolutionary history studies of CLL, these two genetic lesions tend to be founding clonal events while *SF3B1* mutations are often subclonal events in the evolution of CLL^16,45^. Because of this, *SF3B1* mutation is often suggested to promote CLL progression rather than initiation^11^. In rare cases, CLL may progress to a type of B cell lymphoma called Richter syndrome that is characterized by notably higher levels of copy number variations (CNVs) compared to the antecedent CLL^41,46,47^. Interestingly, these CNV events often exist downstream of *SF3B1*-mutated Richter transformed CLL, suggesting that *SF3B1* mutation drives clonal evolution via CNV changes^41,46,47^. Large-scale clonal hematopoiesis studies have also provided supporting evidence for *SF3B1* mutation and CNV changes in potentiating oncogenesis^48,49^. Therefore, we hypothesize that *SF3B1* mutation-driven CNV changes may serve as a general mechanism contributing to clonal evolution.

We first used a tool called RNAseqCNV^36^ to infer the CNV status in our *SF3B1* WT and mutant cell lines. Across all cell lines, we were able to detect a few significant CNV events associated with *SF3B1* mutation, hinting that *SF3B1* mutation may play a conserved role in driving the acquisition of these events (Supplementary Fig. 8a-c). Supported by these initial observations, we next carried out whole genome sequencing (WGS) of all *SF3B1* WT and mutant cell lines. Consistent with our RNAseqCNV results, WGS revealed widespread genetic deletions and amplifications that tend to localize near telomeric and centromeric regions (**Fig4.a-c**). Through WGS, we consistently found significant CNV events identified through RNAseqCNV such as a deletion of chromosome 4p in Nalm-6 (**Fig. 4a** and Supplementary Fig. 8a). HG-3 contained high prevalence of focal, rather than arm level, genetic deletions and amplifications that were invisible to RNAseqCNV but detectable through high resolution WGS (**Fig. 4b** and Supplementary Fig. 8b). In contrast, MEC-1 was characterized more by large-scale genetic changes at the chromosomal arm level (**Fig. 4c** and Supplementary Fig. 8c). Higher level of CNV events also appear to be associated with *SF3B1* mutation in primary CLL patient samples with either *del(13q)* or *del(17p)*, the two most common genetic lesions in CLL^16,41,44^ (**Fig. 4d**), suggesting that CNV changes are a general consequence of *SF3B1* mutation.

We next sought to determine what clinically relevant CNVs, if any, are shared between our cell lines and primary CLL patient samples. In the HG-3 cell line that contains the pre-existing *del(13q)* genetic lesion, we identified genetic deletions at chromosomal locations 17q25.1, 1p31.3, 1p36.21b, and 9q34.3 that are shared with CLL patient samples containing both *SF3B1* mutation and *del(13q)* (**Fig. 4e**). In the MEC-1 cell line that contains the pre-existing *del(17p)* genetic lesion, amplification at chromosome 7q22.1 stands as the most significant CNV event shared with primary CLL patients harboring both *SF3B1* mutation and *del(17p)* (**Fig. 4f**). Nalm-6, in contrast, shared almost no common CNV events with CLL. These results demonstrate that the CLL *SF3B1* mutant cell lines recapitulate CLL-specific CNVs. To confirm this further, we focused on amplification of chromosome 7q found in our MEC-1 cell line. Amplifications on chromosome 7 are frequent CNV events in CLL and a common downstream CNV event of *SF3B1* mutation following transformation of CLL to Richter Syndrome^16,47^. Breaking down WGS and RNAseqCNV calls for chromosome 7 in MEC-1, we confirmed arm level amplification of chromosome 7q starting at q22.1 (**Fig. 4g** and Supplementary Fig. 8d). These results collectively show that general and lineage-specific CNV events are hallmarks of *SF3B1* mutation that drive clonal evolution. Taken together, we have demonstrated that PRECIS engineering can result in disease-relevant *SF3B1* mutant cell line models, providing a powerful tool for further investigation into *SF3B1* mutation in different cancers.

## Discussion

Despite *SF3B1* being the most recurrently mutated splicing gene in cancers, there exists remarkably few isogenic cell lines modeling for *SF3B1* mutation in the disease contexts. Difficulties in engineering *SF3B1* have impeded further research into the roles of oncogenic mutations on this critical splicing factor. In this study, we established a novel, precise, and highly efficient method for introducing *SF3B1* mutation at the endogenous gene locus across multiple cellular contexts based on prime editing technology. This proof-of-concept approach not only provides insights for isogenic cell line engineering and drug development, but also reveals novel aspects of cancer biology.

First, our all-in-one prime editing approach outperforms conventional Cas9 and AAV template-mediated editing as well as base editing technology. We demonstrated that this system can introduce K700E mutation cell lines that model MDS (MDS-L), AML (MOLM-13), and CLL (HG-3 and MEC-1). While our approach uses plasmids for cell line engineering, delivery of the prime editing system using mRNAs and synthetic sgRNAs may potentially simplify and improve editing especially in hard-to-edit cell types where electroporation of large cargos can be tricky. The use of ribonucleoproteins (RNPs) and even viral platforms as delivery methods may even allow for highly efficient *SF3B1* engineering in primary cells such as CD34+ HSPCs, expanding the study of *SF3B1* mutation into primary human contexts. With the recent establishment of a prime editing PE2 murine model, future *in vivo* prime edited models of mutant *SF3B1* and even other splicing factors can also be rapidly established^50^.

Second, the use of synthetic intron reporter systems marking *SF3B1* mutant versus WT cells offers opportunities for further genetic engineering and screening. In this paper, we showed that selection of edited cells with the K700E reporter significantly improved the efficiency for establishment of *SF3B1* mutant cell lines. As the mutation responsiveness of the *MTERF2* synthetic intron is not limited to the K700E mutation in *SF3B1*, engineering possibilities may also be expanded to include other hotspot mutations such as the H662Q mutation (Supplementary Fig. 3h)^30^. With future discovery of additional mutation-responsive introns, prime editing may also be used to engineer hotspot mutations on other splicing factors such *SRSF2*, *U2AF1*, *ZRSR2*, genes encoding the U1 snRNA, and *HNRNPH1*^13^. Because the K700E reporter effectively marks mutant cells from WT cells via GFP expression, this reporter can potentially be paired with a high-throughput CRISPR screening and FACS sorting to identify cofactors essential for *SF3B1* mutation-dependent aberrant splicing.

Third, our method could play a critical role in future development of SF3B1 mutation splicing inhibitor. In the past decade, splicing inhibitors have been developed to directly target SF3B1 such as pladienolides B and its later derivatives^51,52^. However, effective drugging of mutant SF3B1 remains elusive owing to the poor differential sensitivity between *SF3B1* mutated and unmutated cancers and high level of cytotoxicities^53,54^. One possible explanation is presence or absence of additional cofactors that coordinate splicing with mutant spliceosome; For example, mutation on SF3B1 attenuates recruitment of SUGP1, a G-PATCH protein, that recruits DHX15 to surveil and quality control splicing under suboptimal conditions thereby potentiating cryptic splicing^55,56^. Advances in cryo-EM technologies have started to elucidate the dynamics of the early splicing where faithful branchpoint selection is made through several high-resolution structures of early spliceosome^57–60^. However, to date, no structure of mutant spliceosome has been derived, with cancer-associated hotspot mutations only being modeled *in silico*, thereby leaving a resolution gap. For cryo-EM studies, spliceosome particles are purified through *in vitro* isolation from mammalian nuclear extracts that are commonly derived from adherent cell lines such as HeLa and HEK293T. In this paper, we showed that *SF3B1* mutation can be easily engineered into HEK293T without necessitating the K700E reporter or lengthy cloning steps. Thus, prime editing serves as a powerful and effective method for the rapid and facile generation of *SF3B1* mutant HEK293T cell lines that can be used to prepare mutant spliceosomes for cryo-EM studies, to elucidate novel SF3B1 interaction partners and dependencies, and for screening of putative drug candidates.

Finally, our study on disease-related *SF3B1* CLL cell lines identified CLL *SF3B1* mutation-specific alternative RNA splicing as well as revealed the roles of *SF3B1* mutation in inducing CNV changes to drive clonal evolution. The use of the PRECIS methodology in future studies can be set up in animal models and even primary human tissues to study this dynamic in detail to establish a causal relationship. Moreover, the simplicity of *SF3B1* engineering with prime editing offers benefits for drug development and validation. The preponderance of *SF3B1* mutation towards driving genome-wide CNV changes can act as actionable therapeutic vulnerabilities to be exploited to kill *SF3B1*-mutated cancers^14,61^. Our preliminary findings hint at generally conserved mechanism underpinning the acquisition of cancer-associated CNVs that is predicated on the presence of specific pre-existing genetic lesions^62^. The types of CNVs acquired can potentially regulate the clonal trajectory and evolution of *SF3B1*-mutated cancers and may serve to inform therapeutic responses to different inhibitors. Expansion of a cohort of *SF3B1* mutant cell lines with a wider range of genetic contexts would allow for more accurate pharmacological screening and aid in precision medicine.

Collectively, we demonstrated a proof-of-concept approach that is a reliable, efficient, and cost-effective way to engineer *SF3B1* K700E mutation into diverse cell lines with broad implications for cancer research and drug development.

---

**Supplementary Figure 1:**
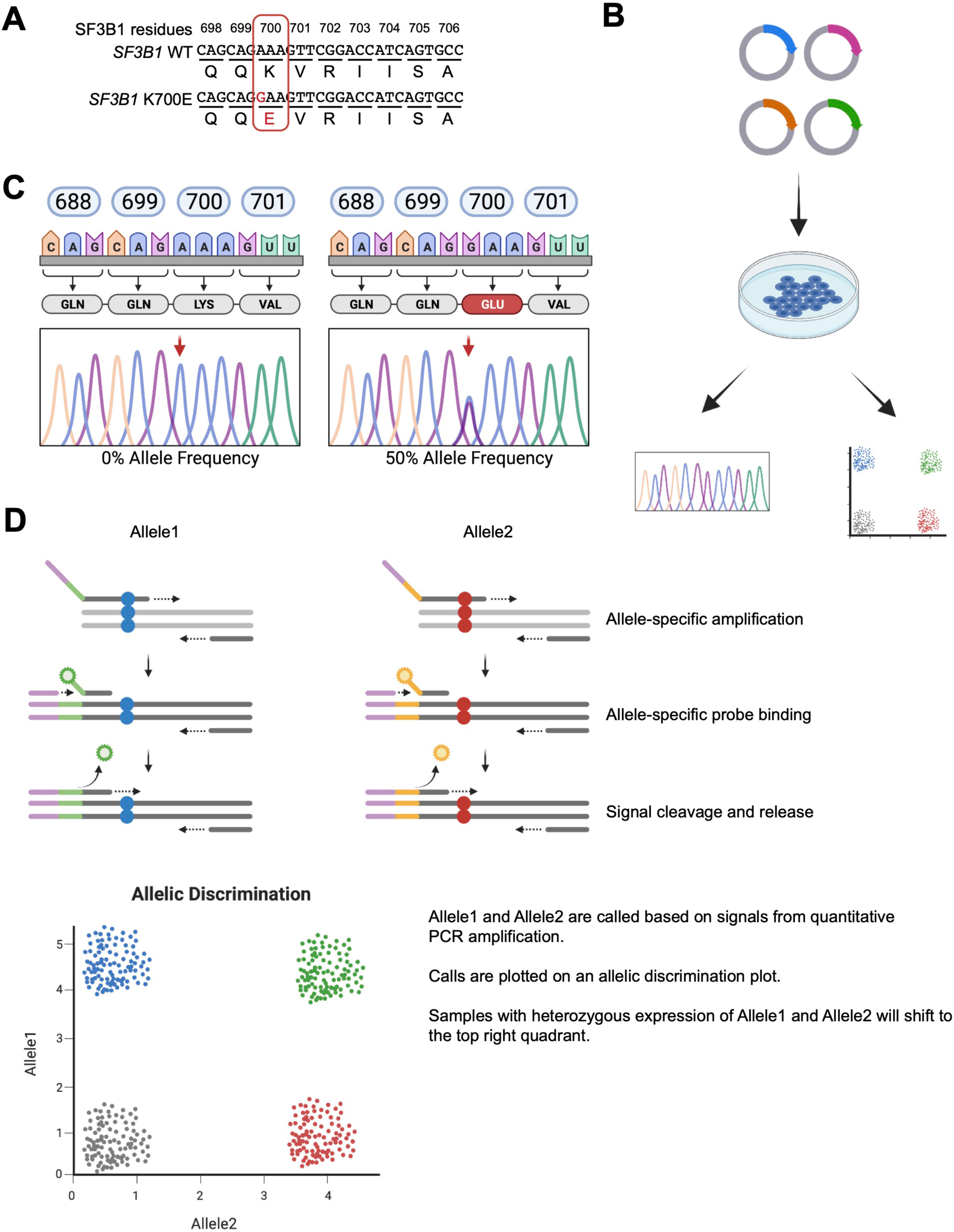
Overview of the K700 locus and mutation validation strategies. A) The *SF3B1* K700 amino acid residue is encoded by a tri-nucleotide AAA sequence. Conversion of the first A>G will result in the hotspot K700E mutation. B) Workflow for testing prime editing of the K700E mutation. After transfecting, the prime editing components into HEK293T, the K700E editing efficiency is assessed via Sanger sequencing and rhAMP SNP assay. C) Cartoon representation of the expected Sanger sequencing trace for the K700 locus. The *SF3B1* K700E mutation is always heterozygous, reaching at most 50% allele frequency in diploid cells. D) Overview of the rhAMP SNP assay; Amplification by allele specific primers results in release of fluorescent signals that can be mapped on an allelic discrimination plot to give a readout on the presence of a SNP in each sample.

**Supplementary Figure 2:**
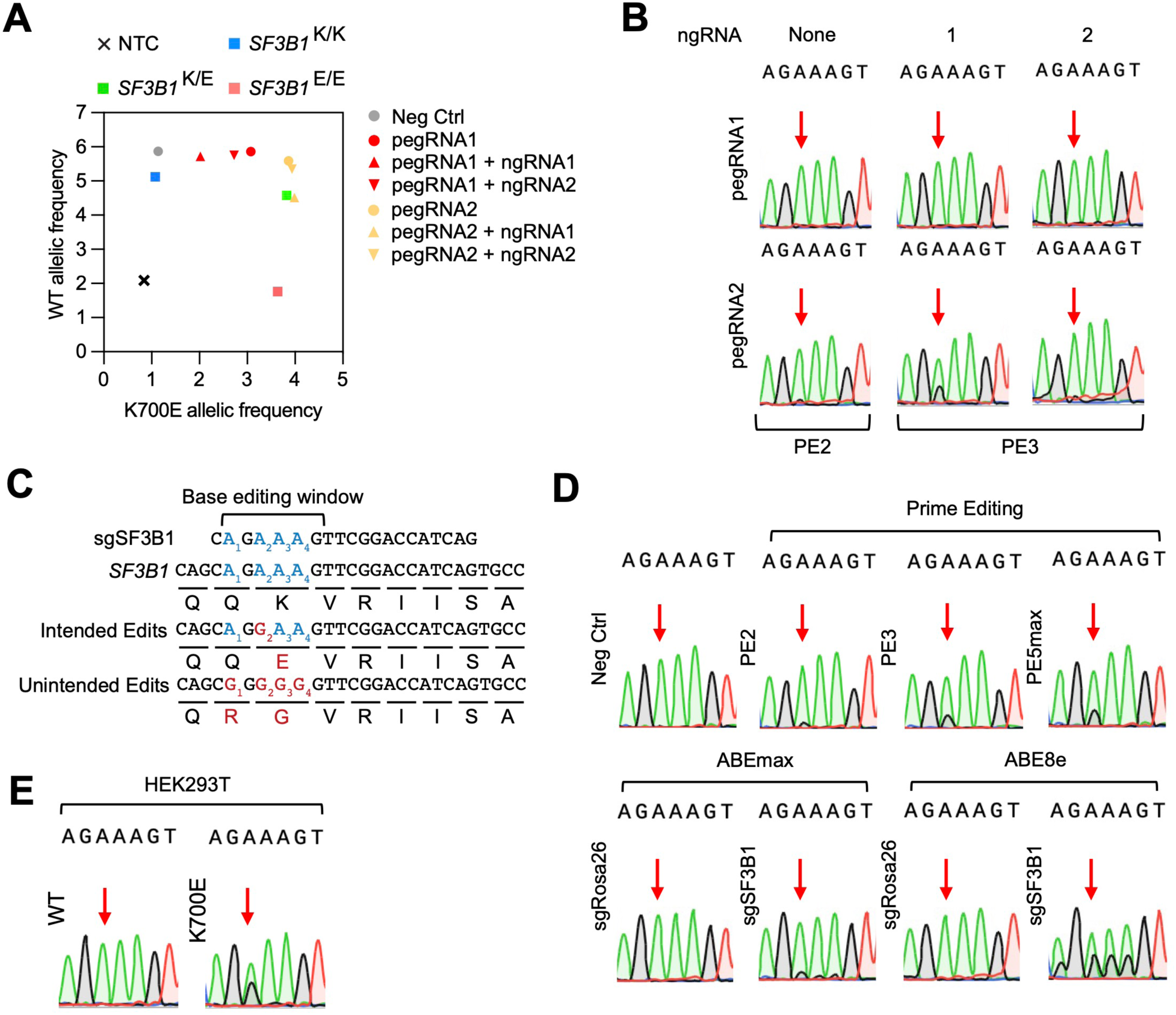
PE2 and PE3 outperform base editing in engineering the K700E mutation. A) Allelic discrimination plot showing prime editing of the K700E mutation using different pegRNAs alone or in combination with ngRNAs based on rhAMP SNP assay. B) Sanger sequencing results for HEK293T transfected with pegRNAs (PE2) alone or in combination with ngRNAs (PE3). C) Base editors are guided to the K700 locus via sgRNA to effectuate A>G editing within the base editing window. Both intended desired editing and unintended collateral editing are shown. D) Sanger sequencing results for prime editing versus base editing in HEK293T at the K700 locus. E) Sanger sequencing results for isogenic HEK293T *SF3B1* WT and K700E clones.

**Supplementary Figure 3:**
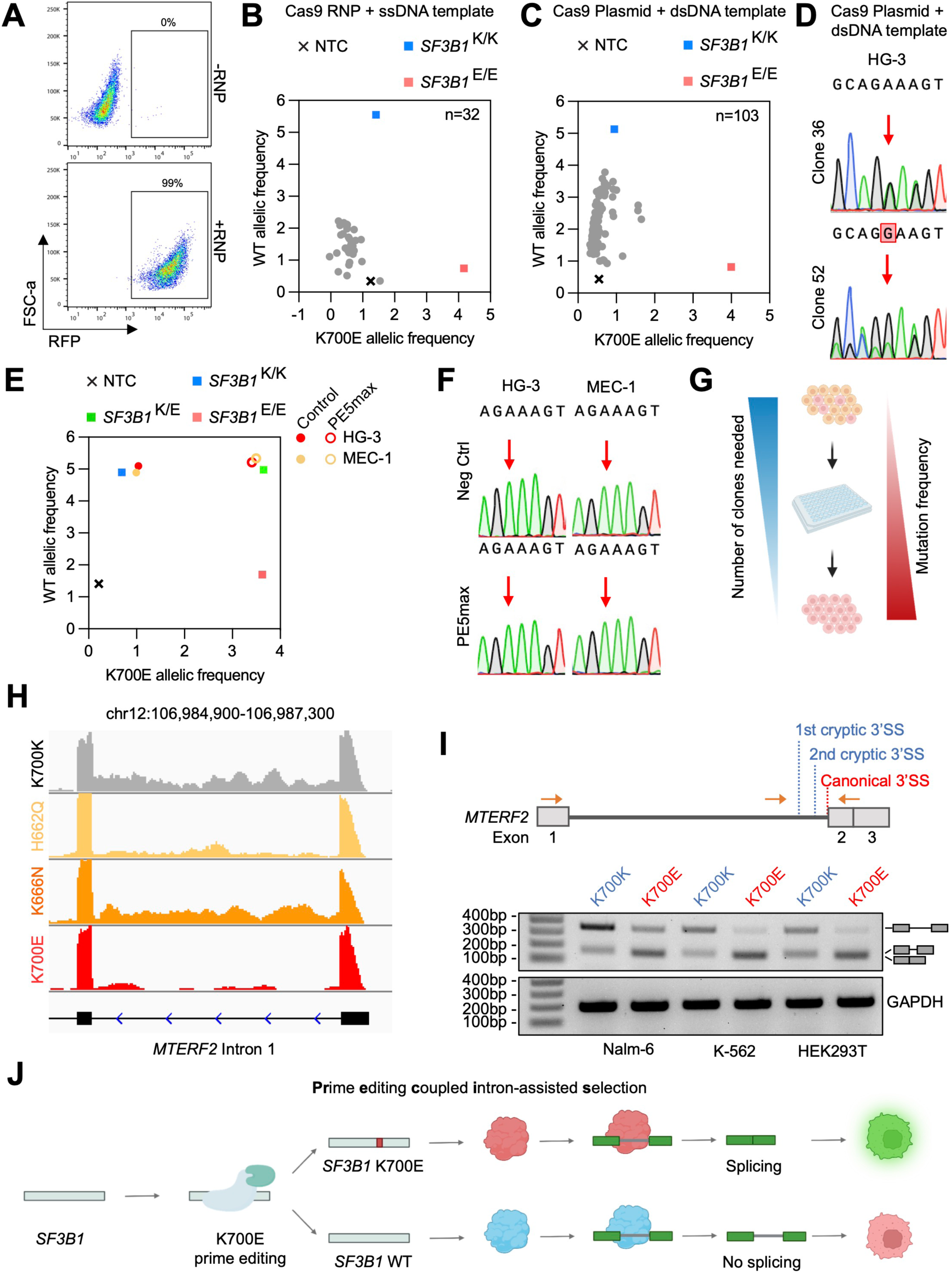
Engineering *SF3B1* mutation into CLL cell lines necessitates an additional enrichment marker. A) Flow cytometry plots showing the electroporation of Cas9 RNP and ssDNA into HG-3. Color indicator is provided by ATTO-550 conjugated to the tracRNA. Allelic discrimination plot for single cell clones of HG-3 cells electroporated with B) Cas9 RNP and ssDNA or C) Cas9-GFP plasmid and dsDNA repair template. GFP is used as an indicator to sort for cells electroporated with plasmids before single cell cloning. D) Sanger sequencing results for two single cell clones isolated from HG-3 cells electroporated with Cas9-GFP plasmid and dsDNA repair template. E) Allelic discrimination plot and F) Sanger sequencing for prime editing by PE5max K700E in HG-3 and MEC-1 cells. G) For single cell cloning, higher efficiency editing in bulk cell populations will result in less clones needing to be screened to isolate pure *SF3B1* K700E clones. H) Nalm-6 RNA-seq reads for the *MTERF2* intron 1 splicing between isogenic WT and different *SF3B1* mutant clones. I) (top) Primer designs and (bottom) PCR for checking the splicing status on *MTERF2* intron 1 in different *SF3B1* WT and K700E cell lines. J) Overview of the PRECIS workflow: After the PE5max K700E is used to introduce the K700E mutation into cells, mutant *SF3B1* will splice the K700E reporter to give GFP expression as a marker for prime edited cells.

**Supplementary Figure 4:**
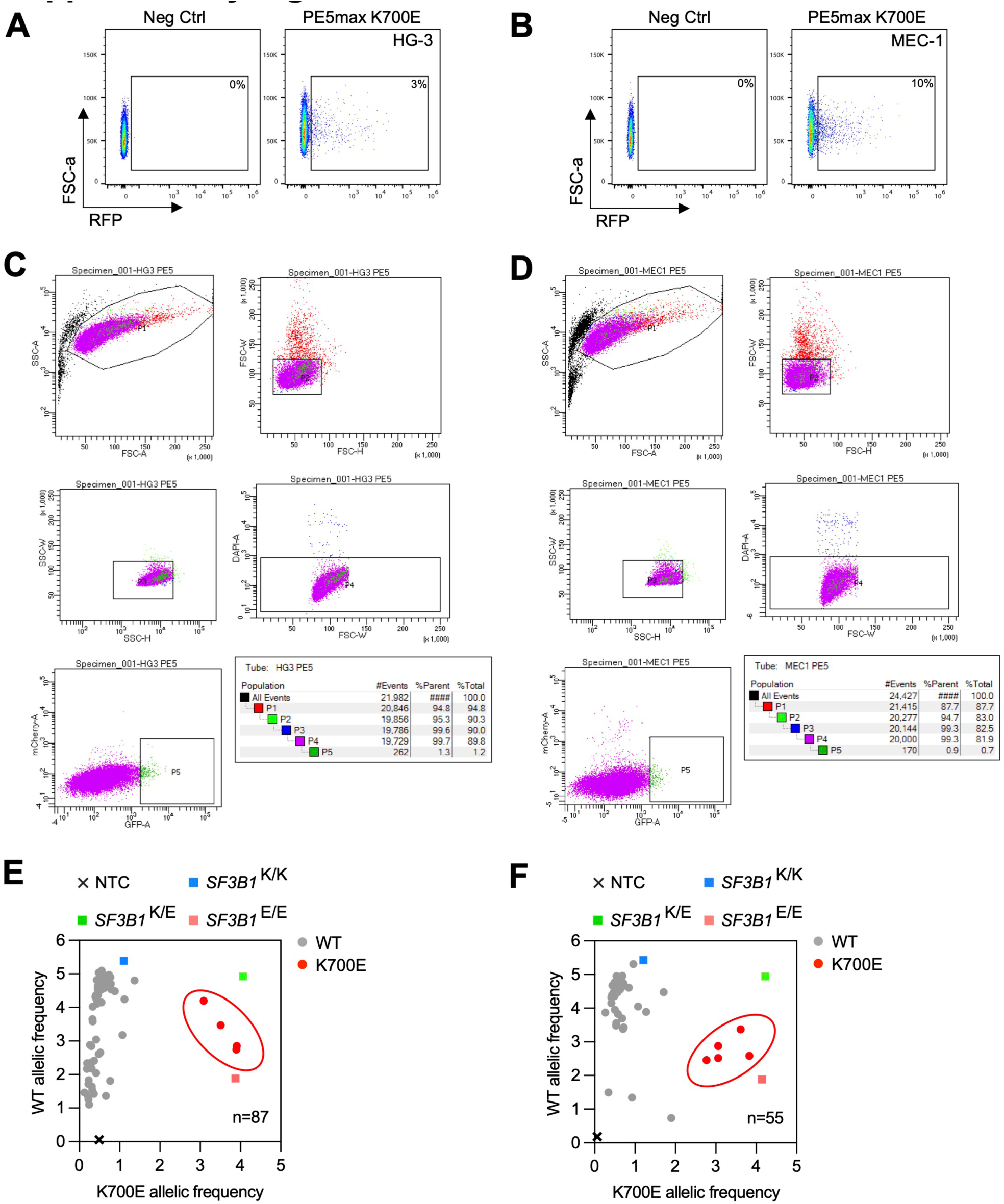
FACS sorting and RhAMP SNP screening yield isogenic *SF3B1* mutant clones. Flow cytometry plots showing the PE5max K700E electroporation efficiency and population targeted for sorting for A) HG-3 and B) MEC-1. Workflows for enriching the GFP bright population in C) HG-3 and D) MEC-1 cells sorted for PE5max K700E. Allelic discrimination plots showing the rhAMP SNP screening of single cell clones of E) HG-3 and F) MEC-1 cells sorted for GFP bright cells.

**Supplementary Figure 5:**
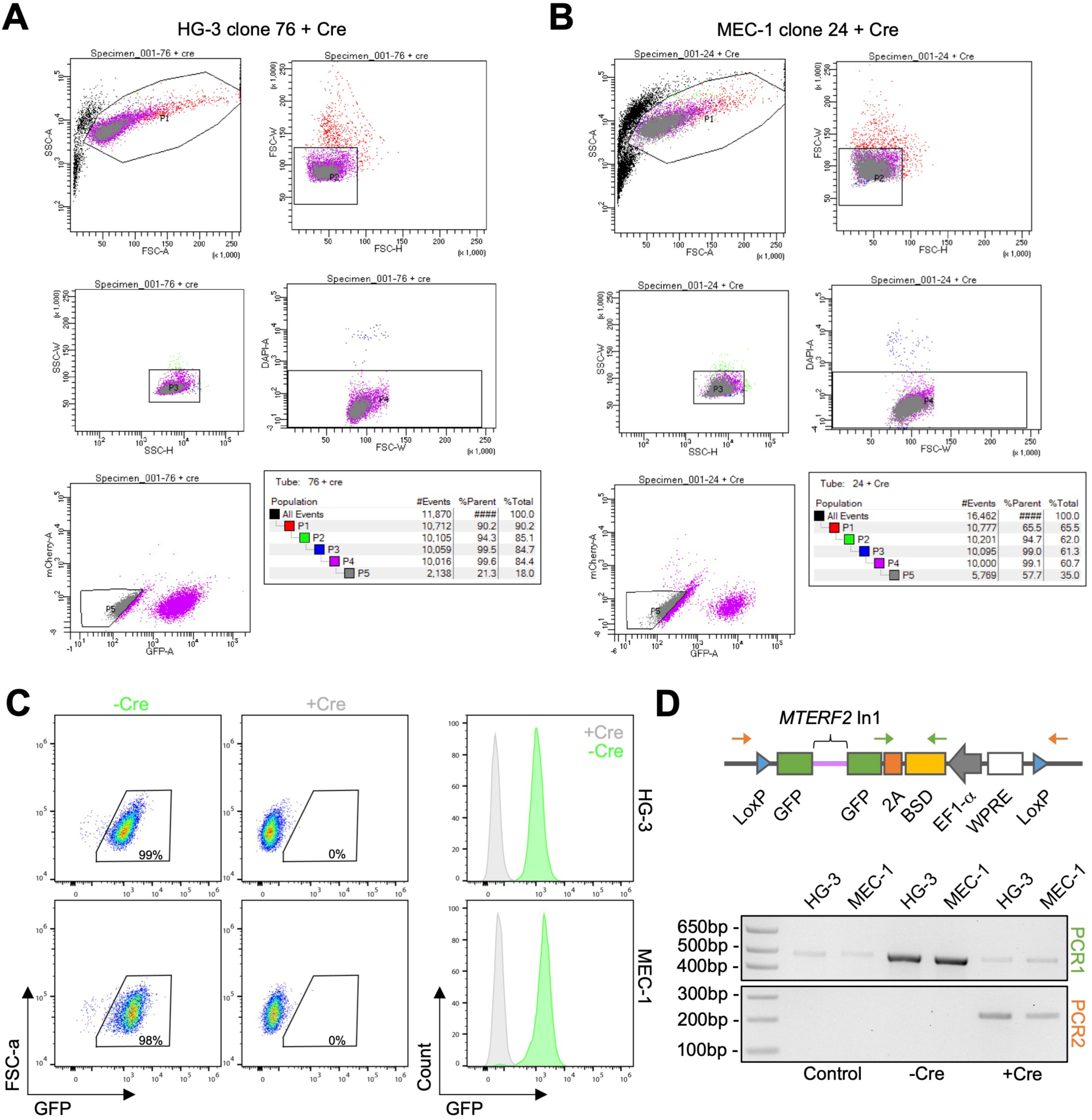
The K700E reporter is removed via Cre-mediated recombinase treatment. Workflows for sorting out GFP negative cells following Cre overexpression for A) HG-3 and B) MEC-1 *SF3B1* K700E cells. C) Flow cytometry plots (left) and histograms (right) showing GFP expression with and without Cre overexpression in HG-3 and MEC-1 *SF3B1* K700E cells. D) (top) Primer designs and (bottom) PCR to genotype for deletion of the K700E reporter.

**Supplementary Figure 6:**
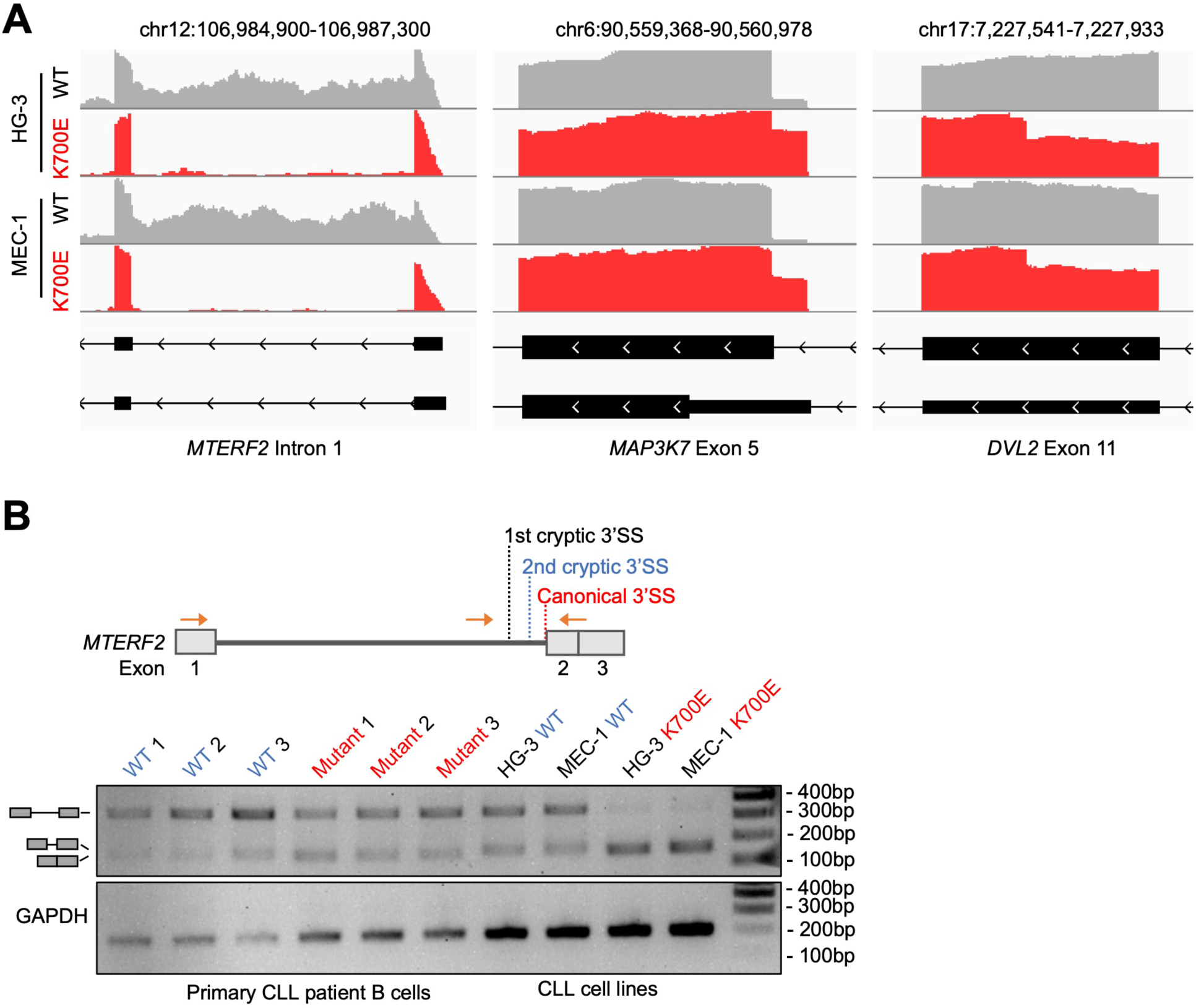
Splicing analysis of HG-3 and MEC-1 *SF3B1* mutant cell lines. A) RNA-seq analyses showing alternative splicing on *MTERF2*, *MAP3K7*, and *DVL2* between HG-3 and MEC-1 *SF3B1* WT and K700E cells. B) (top) Primer designs and (bottom) PCR for checking *MTERF2* intron 1 splicing in primary CLL patient samples versus CLL cell lines.

**Supplementary Figure 7:**
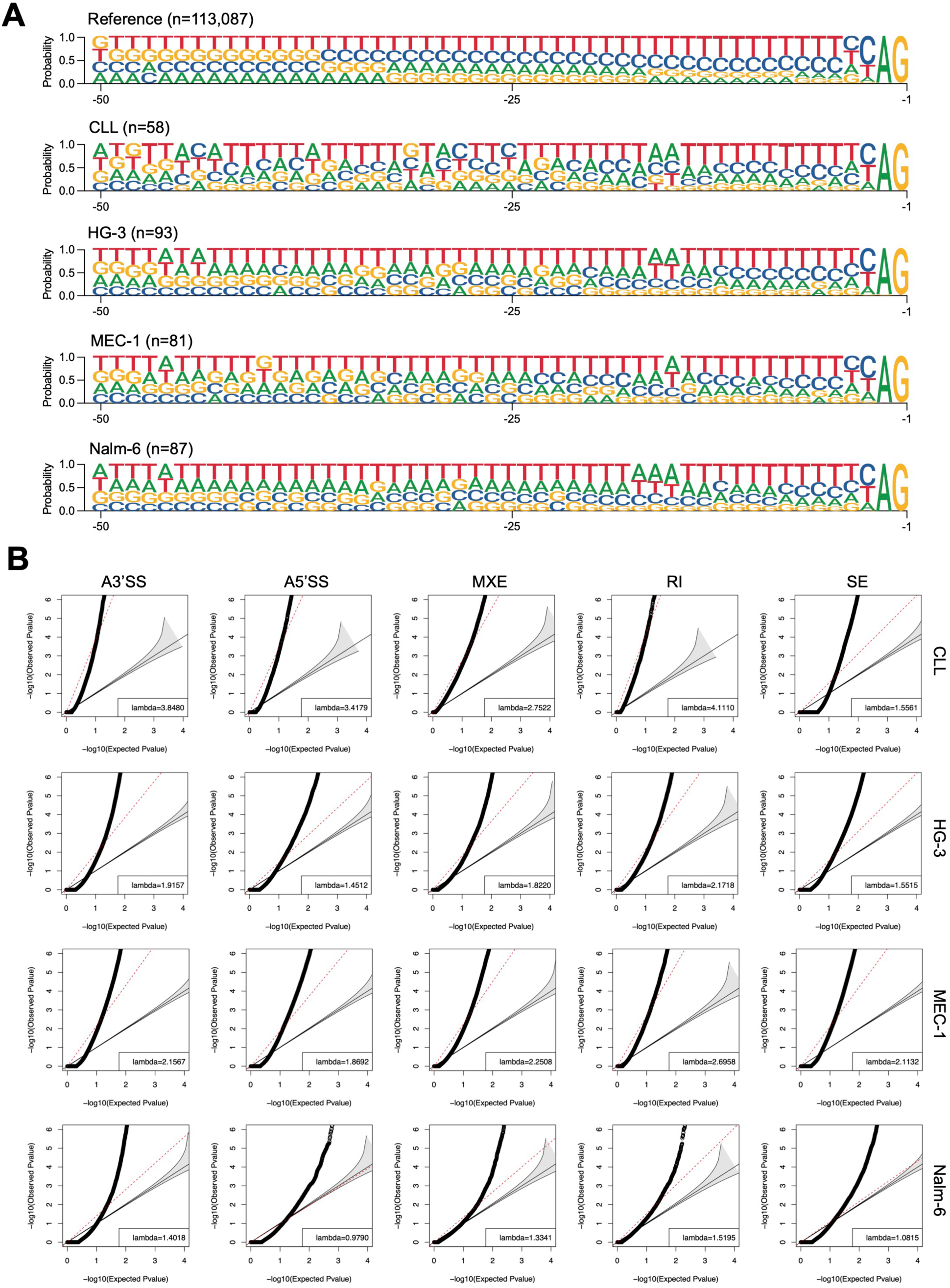
Aberrant 3’SS splicing profiles of *SF3B1*-mutated cell lines and primary CLL. A) Logo plots showing the cryptic 3’SS sequence and location in *SF3B1*-mutated primary CLL samples and cell lines versus the RefSeq reference. B) Q-Q plots of observed P values versus expected P values for five types of alternative splicing events in *SF3B1*-mutated primary CLL samples and cell lines. Least-squares linear fit with slope γ for the lower 95^th^ percentile is indicated by red lines. 95% confidence intervals are represented by the gray-shaded area.

**Supplementary Figure 8:**
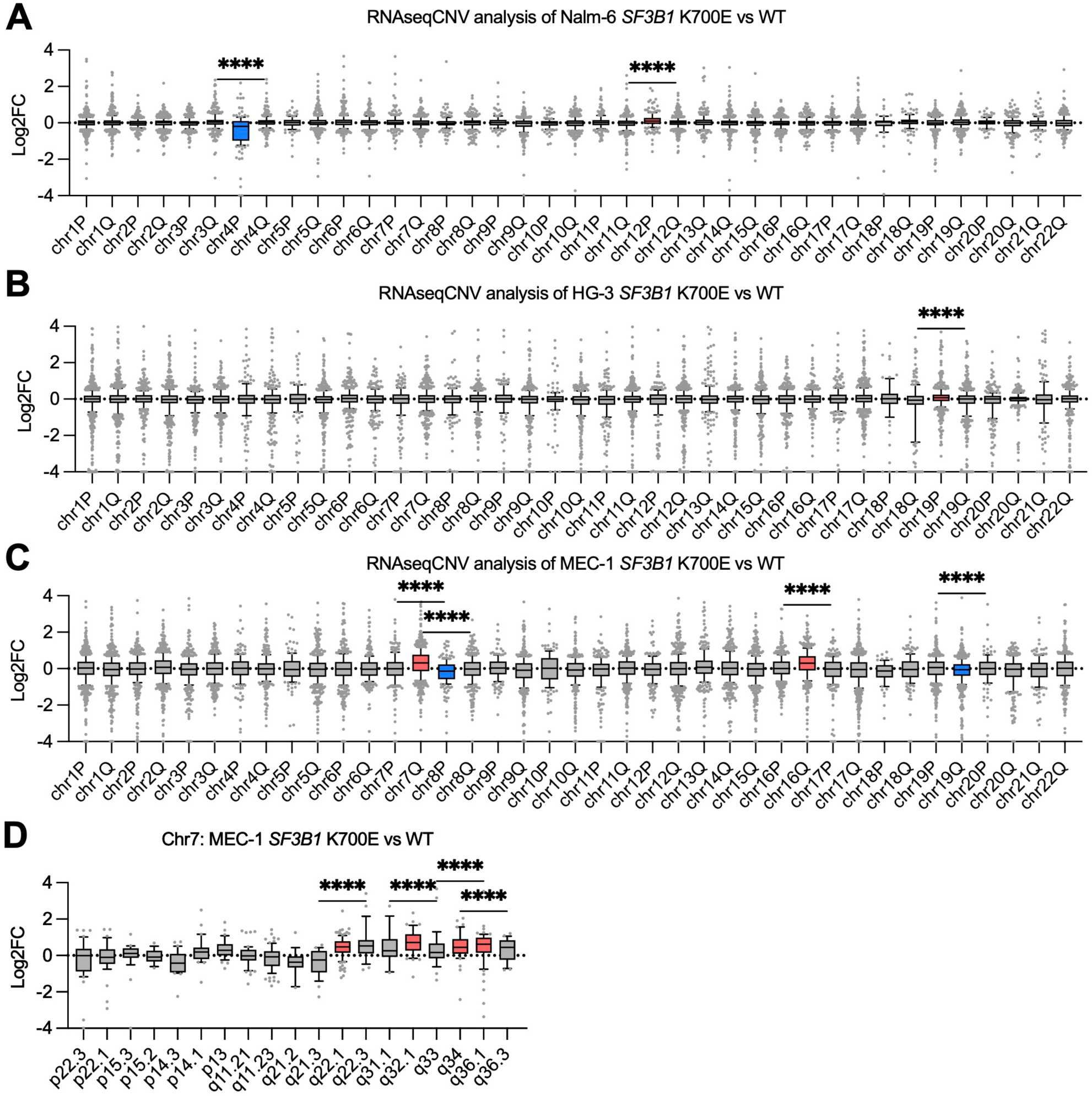
RNAseqCNV analysis uncovers widespread CNV events in *SF3B1* mutant cell lines. RNAseqCNV analysis represented by bar plot based on gene expressions at each indicated chromosomal loci for A) Nalm-6, B) HG-3, and C) MEC-1. Regions that underwent amplification and deletion are shown in red and blue, respectively. Only chromosomal locations that have passed the one sample Wilcoxon signed rank test with a high statistical cutoff (****P≤0.0001) were annotated for deletion or amplification. D) Bar plot of gene expressions on chromosome 7. A chi-squared test was performed with -log_10_(0.05) as the statistical cutoff.

**Supplementary Table 1:** List of current SF3B1 mutant cell lines.

| Supplementary Table 1: List of current *SF3B1* mutant cell lines |  |  |  |  |
| --- | --- | --- | --- | --- |
| Cell Line | Method | Mutation | Disease | Source |
| HEK293T | Cas9 HDR | K666T | Human Kidney Fibroblast | Alsafadi et al, *Nature Communications* 2016 |
| HEK293T | Cas9 HDR | K700E | Human Kidney Fibroblast | Cusan et al, *Journal of Clinical Investigation* 2023 |
| MCF-10A | Cas9 HDR | K700E | Breast Cancer | Liu et al, *Journal of Clinical Investigation* 2020 |
| MCF-10A | AAV HDR | K700E,<br>R702R | Breast Cancer | Dalton et al, *Journal of Clinical Investigation* 2019 |
| hTERT-IMEC | AAV HDR | K700E,<br>R702R | Mammary Epithelial Cells | Dalton et al, *Journal of Clinical Investigation* 2019 |
| CCE mESC | Cas9 HDR | K700E | Mouse Embryonic Stem Cells | Gupta et al, *Nucleic Acids Research* 2018 |
| B16F10 | Cas9 HDR | R1074H | Mouse Melanoma Cells | Chang et al, *Journal of Biological Chemistry* 2021 |
| K-562 | AAV HDR | K700E | Chronic Myelogenous Leukemia | Dharman et al, *Cell Reports* 2015 |
| K-562 | Cas9 HDR | K700E | Chronic Myelogenous Leukemia | Liberante et al, *Scientific Reports* 2019 |
| K-562 | AAV+Cas9 HDR | K700E | Chronic Myelogenous Leukemia | Boddu et al, *Communication Biology* 2021 |
| K-562 | Cas9 HDR | K700E | Chronic Myelogenous Leukemia | Mian et al, *Science Translational Medicine* 2023 |
| Nalm-6 | AAV HDR | H662Q,<br>K666N,<br>K700E | Acute Lymphoblastic Leukemia | Dharman et al, *Cell Reports* 2015 |
| MEC-1 | Cas9 HDR | K700E | Chronic Lymphocytic Leukemia | Lopez-Oreja et al, *Life Science Alliance* 2023 |

